# Human and rat skeletal muscle single-nuclei multi-omic integrative analyses nominate causal cell types, regulatory elements, and SNPs for complex traits

**DOI:** 10.1101/2020.07.01.183004

**Authors:** Peter Orchard, Nandini Manickam, Arushi Varshney, Vivek Rai, Jeremy Kaplan, Claudia Lalancette, Katherine Gallagher, Charles F. Burant, Stephen C.J. Parker

## Abstract

**Background:** Skeletal muscle accounts for the largest proportion of human body mass, on average, and is a key tissue in complex diseases, mobility, and quality of life. It is composed of several different cell and muscle fiber types.

**Results:** Here, we optimize single-nucleus ATAC-seq (snATAC-seq) to map skeletal muscle cell-specific chromatin accessibility landscapes in frozen human and rat samples, and single-nucleus RNA-seq (snRNA-seq) to map cell-specific transcriptomes in human. We capture type I and type II muscle fiber signatures, which are generally missed by existing single-cell RNA-seq methods. We perform cross-modality and cross-species integrative analyses on 30,531 nuclei, representing 11 libraries, profiled in this study, and identify seven distinct cell types ranging in abundance from 63% (type II fibers) to 0.9% (muscle satellite cells) of all nuclei. We introduce a regression-based approach to infer cell types by comparing transcription start site-distal ATAC-seq peaks to reference enhancer maps and show consistency with RNA-based marker gene cell type assignments. We find heterogeneity in enrichment of genetic variants linked to complex phenotypes from the UK Biobank and diabetes genome wide association studies in cell-specific ATAC-seq peaks, with the most striking enrichment patterns in muscle mesenchymal stem cells (∼3% of nuclei). Finally, we overlay these chromatin accessibility maps on GWAS data to nominate causal cell types, SNPs, and transcription factor motifs for creatinine levels and type 2 diabetes signals.

**Conclusions:** These chromatin accessibility profiles for human and rat skeletal muscle cell types are a useful resource for investigating specific cell types and nominating causal GWAS SNPs and cell types.

## Background

Skeletal muscle tissue accounts for 30-40% of body mass, which is the largest tissue, on average, in adult humans and is central to basic quality of life and complex diseases (1,2). Like other tissues, skeletal muscle is composed of a mixture of different cell types. Most of the tissue is composed of muscle fibers, which may be categorized into different fiber types, each of which display distinct metabolic and molecular phenotypes. The proportion of muscle fibers accounted for by each fiber type varies across individuals (3). Muscle-related diseases may differentially impact different fiber types, and fiber type proportions are associated with complex phenotypes, including aerobic and anaerobic exercise capacity and type 2 diabetes (T2D) status (4). Muscle satellite cells are progenitors to muscle fibers, indispensable for the generation and regeneration of muscle (5); these cells are present in skeletal muscle tissue, as are several other cell types, such as mesenchymal stem cells, that cooperate in muscle regeneration (6,7). Molecular associations with skeletal muscle tissue/muscle fiber characteristics and muscle-related complex diseases could be mediated in part by these stem cell-like populations; for example a genetic variant that alters the developmental of a satellite cell could carry important implications for later muscle function, just as some T2D-associated variants are proposed to impact pancreatic/beta cell development rather than the function of mature beta cells (8,9) and facial morphology associated variants may act through progenitor cell populations (10). Immune cells infiltrate muscle tissue and communicate with muscle cells as well, playing a particularly important role following injury (11). Profiling the transcriptomic and epigenomic landscapes of these cell types and muscle fiber types may therefore contribute to our understanding of the biology of muscle development and muscle-related complex traits.

Bulk profiling of skeletal muscle tissue ignores this heterogeneity and is dominated by the most common cell types (muscle fibers), but single-cell/-nucleus methods overcome this and allow profiling of the constituent cell types. In the case of skeletal muscle, the distinction between single-nucleus and single-cell profiling is particularly important as (1) skeletal muscle fibers have an elongated shape that may make them difficult to capture in single-cell suspensions, and (2) muscle fibers are multinucleated, meaning that a single-cell measurement will capture the output of many nuclei. Previous single-cell RNA-seq studies of human (12–14), mouse (15–20), and pig (21) skeletal muscle tissue either capture no muscle fiber nuclei or capture them in unrepresentative proportions. Bulk analysis of pooled, dissected muscle fibers have generated fiber-type specific transcriptional profiles (22–25) and analysis of specific isolated muscle resident cell populations (26–28) have generated insights into targeted cell subpopulations but these studies are necessarily biased towards specific cell types. To date no single nucleus ATAC-seq (snATAC-seq) studies of whole human or rat skeletal muscle tissue samples has been performed.

Here, we employ single-nucleus RNA-sequencing (snRNA-seq) and ATAC-seq (snATAC-seq) on the 10X Genomics platform to profile gene expression and chromatin accessibility of frozen skeletal muscle cell populations in human and rat. First we examine the influence of fluorescence activated nucleus sorting (FANS) and nucleus loading concentration on the performance of the platform. Next, we perform joint clustering of the snRNA-seq and snATAC-seq libraries to determine the cell types detected in skeletal muscle tissue samples and map their respective transcriptomes and chromatin landscapes. We then integrate the resulting genomic maps with UK Biobank and T2D-related GWAS results to explore the relationship between these cell types and a broad range of human phenotypes and diseases and nominate causal SNPs at several genomic loci.

## Results

### FANS negatively impacts 10X snATAC-seq results

Before being loaded onto the 10X platform, nuclei must be isolated from the samples of interest. This process involves cell lysis, which produces viable nuclei as well as substantial cellular debris and dead nuclei, some of which inevitably remains in the final nuclei suspension. By staining the DNA in live nuclei and using FANS to selectively filter the suspension for stained entities, one should be able to remove dead nuclei and cellular debris in the suspension, improving the purity and quality of the suspension loaded onto the 10X platform. However, the FANS process could stress the nuclei or otherwise alter the snRNA-seq and snATAC-seq results. Comparing quality control metrics and (in the case of snRNA-seq) aggregate gene expression or (in the case of snATAC-seq) aggregate ATAC-seq peaks/signal between snRNA-seq and snATAC-seq libraries generated from nuclei that either did or did not undergo FANS allows one to detect substantial changes that FANS may introduce. Also, because the aggregate of reads from a snATAC-seq library should resemble the profile of an ATAC-seq library on the same biological sample, one can generate bulk and single-nucleus libraries from a single sample and compare quality control metrics and ATAC-seq signal between them. Therefore, to determine the effect of FANS on 10X snRNA-seq and snATAC-seq results, we performed three nuclear isolations from a single human muscle sample, mixed the resulting nuclei together, and performed FANS (using DRAQ7 staining) on one half of the suspension (Fig. 1A). The FANS and non-FANS suspensions were then each used to produce two replicate snATAC-seq and two replicate snRNA-seq libraries, resulting in eight total libraries (four snATAC and four snRNA). We also generated two independent bulk ATAC-seq libraries from the same biological sample, allowing us to compare snATAC-seq profiles, with and without FANS, to a comparable bulk ATAC-seq profile.

**Figure 1:**
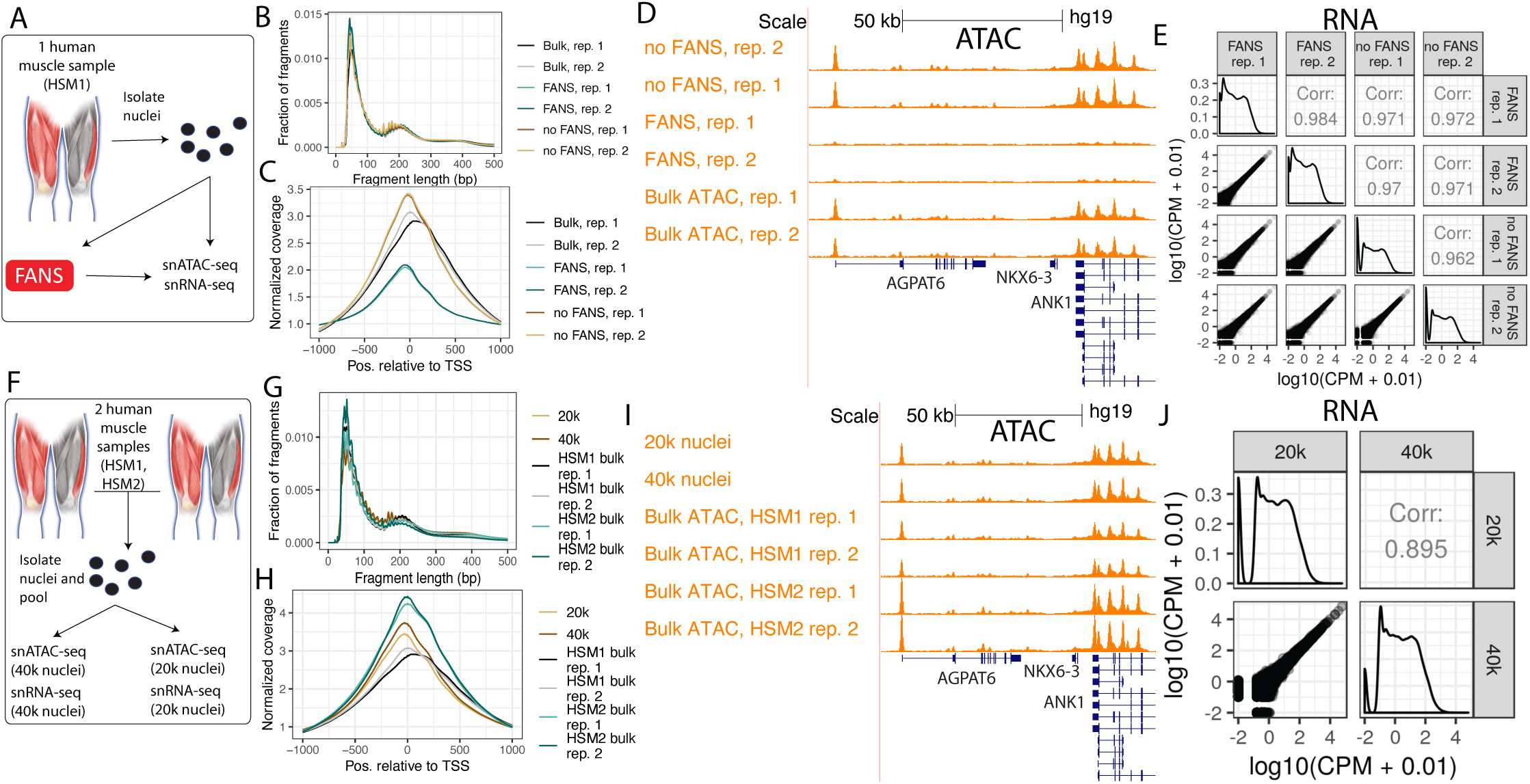
(A) Study design to determine the effect of FANS on snRNA-seq and snATAC-seq results. Muscle cartoon adapted from Scott et al. 2016. HSM1 refers to one specific skeletal muscle sample (‘human skeletal muscle 1’). Bulk ATAC-seq was performed on HSM1 as well (two replicates, each separate nuclei isolations). (B) Fragment length distribution and (C) TSS enrichment for two snATAC-seq libraries that did not undergo FANS and two that did, as well as two bulk ATAC-seq replicates from the same sample (‘Bulk’). (D) ATAC-seq signal at the *ANK1* locus for FANS or non-FANS input snATAC-seq libraries, and the two bulk ATAC-seq libraries. All tracks are normalized to 1M reads. Gene model (GENCODE v19 basic) displays protein coding genes only. (E) Correlation between FANS and non-FANS snRNA-seq libraries; each point represents one gene. (F) Study design to determine the effect of loading 20k vs 40k nuclei into the 10X platform, utilizing HSM1 as well as a second sample, HSM2 (‘human skeletal muscle 2’). Bulk ATAC-seq was performed on HSM1 (same libraries as in (a)) and on HSM2 (two replicates, each separate nuclei isolations). (G) Fragment length distribution and (H) TSS enrichment for snATAC-seq libraries after loading 20k vs 40k nuclei, as well as for the four bulk ATAC-seq libraries (two each from the two muscle samples, ‘HSM1 bulk’ and ‘HSM2 bulk’). (I) ATAC-seq signal at the *ANK1* locus for the 20k and 40k libraries and the four bulk ATAC-seq libraries. All tracks are normalized to 1M reads. Gene model (GENCODE v19 basic) displays protein coding genes only. (J) Correlation between snRNA-seq libraries resulting from loading 20k vs 40k nuclei.

First we examined the four snATAC-seq libraries, comparing the aggregate signal for each library to bulk ATAC-seq libraries from the same biological sample. We called peaks for the four libraries and ran the ataqv quality control software package (29) on the aggregated data to examine the overall transcription start site (TSS) enrichment and fragment length distributions. The fragment length distributions for each library resembled the expected stereotypical ATAC-seq fragment length distribution, showing an abundance of short fragments as well as mononucleosomal fragments (Fig. 1B) (Buenrostro et al., 2013); however, the TSS enrichment was lower in the FANS libraries (Fig. 1C), indicating the FANS libraries had a lower signal to noise ratio. This difference in signal-to-noise ratio is demonstrated when visualizing the ATAC-seq signal at genomic regions active in muscle, such as the *ANK1* locus (Fig. 1D) (30). We additionally overlapped TSS-distal ATAC-seq peaks from each of the libraries with existing chromatin states from diverse tissues and cell types (31) and found that the peaks from the non-FANS libraries showed considerable overlap with skeletal muscle enhancers, while the peaks from the FANS libraries showed poor overlap (Fig. S1). ATAC-seq signal across FANS libraries showed poor correlation with the two bulk ATAC-seq libraries from the same sample (Fig. S2). We therefore concluded that FANS has a clear negative impact on 10X snATAC-seq results.

Next we examined the four snRNA-seq libraries. All four libraries showed high correlation, indicating that FANS does not substantially alter snRNA-seq results, at least at the pseudobulk gene expression level (Fig. 1E). In order to determine if FANS altered the yield of quality nuclei, we used read counts and mitochondrial contamination to select quality nuclei from each library, additionally removing doublets using doubletfinder (Fig. S3) (32). We found that FANS substantially increased the number of quality nuclei obtained (2,004 and 2,078 for non-FANS libraries; 7,715 and 7,118 for FANS libraries). We therefore concluded that FANS has little effect on pseudobulk gene expression measurements, but may alter nucleus yield.

### snATAC-seq and snRNA-seq results are robust to nucleus loading concentrations

The concentration at which nuclei are loaded onto the 10X platform is an important parameter affecting data quality and the number of nuclei available for downstream analysis. Increasing the loading concentration increases the maximum number of nuclei from which data can be obtained; however, it also increases the probability that multiple nuclei end up with the same gel bead, thereby increasing the doublet rate. Balancing these outcomes is important to maximize the amount of quality data and number of nuclei available for downstream analysis. To evaluate the effect of increasing the number of nuclei loaded onto the platform, we performed a separate experiment in which we isolated nuclei from two muscle samples, mixed them together, and then loaded either 20k or 40k nuclei (as quantified by a Countess II FL Automated Cell Counter) into a 10X well for snRNA-seq and for snATAC-seq (Fig. 1F). We also generated two independent bulk ATAC-seq libraries from the biological sample for which bulk ATAC-seq profiles were not already available, allowing us to compare snATAC-seq profiles to comparable bulk ATAC-seq profiles.

The snATAC-seq libraries displayed the expected fragment length distributions and comparable TSS enrichments (Fig. 1G, H). We examined the aggregate signal of the snATAC-seq libraries next to bulk ATAC-seq libraries from the same samples and confirmed that both libraries showed strong signal, comparable to that of bulk data (Fig. 1I). Overlap between TSS-distal ATAC-seq peaks called on both libraries and chromatin states were likewise similar, showing relatively high overlap with skeletal muscle enhancers (Fig. S4), and the ATAC-seq signal in the libraries correlated with bulk ATAC-seq signal to an extent comparable to the correlation between two bulk ATAC-seq libraries (Fig. S5). After selecting quality nuclei (Fig. S6), we found that the higher loading concentration yielded 2,035 nuclei while the lower concentration yielded 855 nuclei (after doublet removal).

Correlation between the snRNA-seq libraries was high, indicating that the loading concentration could be changed substantially without compromising data quality (Fig. 1J). We again found the higher loading concentration yielded more quality nuclei than the lower concentration (3,839 vs 2,118; Fig. S7) after doublet removal.

10X guidelines recommend loading up to 15k nuclei into a well; however, our results indicate that exceeding this loading concentration can still yield quality snATAC-seq results (as measured by standard quality control metrics relative to bulk ATAC-seq data) and, for both snATAC-seq and snRNA-seq, increase the number of quality nuclei even after accounting for the increase in doublet rate. The aggregate gene expression/ATAC-seq signal profile was comparable between loading concentrations. One caveat to these conclusions is that the actual number of nuclei loaded into the well may differ from our estimated numbers, as debris in the nuclei preps may affect the accuracy of the nuclei counts.

### Joint clustering of human and rat snATAC-seq and snRNA-seq identifies skeletal muscle cell types

To determine cell types present in skeletal muscle samples, we selected high-quality ATAC and RNA nuclei from the FANS/non-FANS libraries and the 20k/40k nuclei libraries generated above and performed joint clustering. snATAC-seq libraries that underwent FANS were excluded as they failed to provide quality data. We generated and included a snATAC-seq library containing a mix of human and rat nuclei (Fig. S8, S9; Tables S1, S2). Information about the biological samples and post-QC nucleus summary statistics for each library is provided in Table S3. In total we obtained 24,866 human snRNA-seq (mean UMIs = 7,482), 5,053 human snATAC-seq (mean fragments = 41,655), and 612 rat snATAC-seq (mean fragments = 60,875) nuclei. We used integrative nonnegative matrix factorization (iNMF) as implemented in the LIGER (linked inference of genomic experimental relationships) software package (Welch et al., 2019) to perform joint clustering on snRNA-seq and snATAC-seq nuclei and identified seven cell type clusters (Fig. 2A). Nuclei from different modalities, species, and libraries integrated well, indicating that clustering was not driven by technical factors (Fig. 2B).

**Figure 2:**
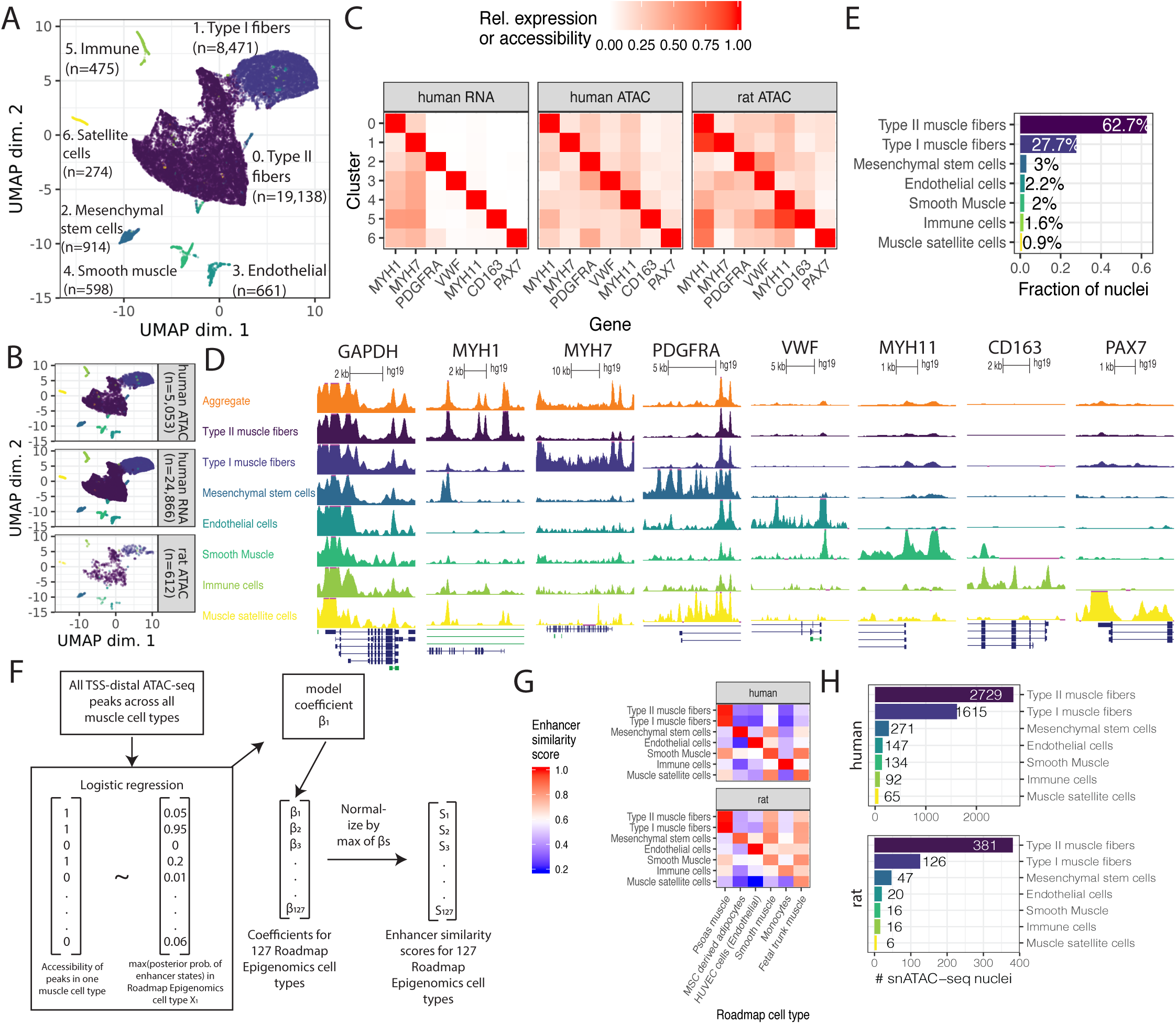
(A) UMAP after clustering human snATAC-seq, human snRNA-seq, and rat snATAC-seq nuclei with LIGER. (B) UMAP facetted by species and modality. (C) Gene expression (snRNA-seq) or accessibility (snATAC-seq; gene promoter + gene body) of marker genes. Values are column-normalized. (D) ATAC-seq signal for human snATAC-seq nuclei in each cluster. All tracks are normalized to 1M reads. (E) Fraction of nuclei, across both species and modalities, assigned to each cell type. (F) Logistic regression-based approach to score similarity between TSS-distal ATAC-seq peaks (> 5 kb from TSS) and Roadmap Epigenomics enhancer states. For all TSS-distal ATAC-seq peaks across all muscle cell types, we scored the accessibility of the peak (0/1) in each of the muscle cell types based on the presence or absence of a peak call in that cell type. Then, for a given one of the 127 Roadmap Epigenomics cell types, we determined the maximum posterior probability of the enhancer states in the Roadmap Epigenomics chromHMM model within each peak. We then used logistic regression to model the relationship between the peak accessibility and the enhancer posteriors (running one model per muscle cell type per Roadmap Epigenomics cell type). Then, for each muscle cell type, the model coefficient was normalized to 1 by dividing by the maximum coefficient across all 127 Roadmap Epigenomics cell types, and this value was used as the enhancer similarity score for that muscle cell type and Roadmap Epigenomics cell type. (G) Similarity of snATAC-seq peak calls for each cell type and species to Roadmap Epigenomics chromHMM enhancer states based on the logistic regression procedure outlined in (F). The Roadmap Epigenomics cell type names have been adjusted slightly for clarity and the sake of space. The full names and the identifiers from the Roadmap Epigenomics paper are: Psoas muscle (E100), Mesenchymal Stem Cell Derived Adipocyte Cultured Cells (E023), HUVEC Umbilical Vein Endothelial Primary Cells (E122), Stomach Smooth Muscle (E111), Primary monocytes from peripheral blood (E029), and Fetal Muscle Trunk (E089). (H) Nucleus counts per species for snATAC-seq data.

We used marker genes to assign cell types to each cluster (Table S4) and found clear concordance between human snRNA-seq and snATAC-seq (Fig. 2C, D). We found marker gene accessibility in the rat snATAC-seq data to be largely consistent with the human data, though examination of the myosin heavy chain genes, often used to distinguish between different muscle fiber types, indicated that a considerable number of rat type II muscle fiber nuclei were likely present in the type I muscle fiber cluster (the opposite did not seem to occur; i.e., the type II muscle fiber cluster appeared to be relatively free of rat type I muscle fiber nuclei; Fig. S10). This mixing of some rat muscle fiber nuclei is a limitation of our data; because only 612 of 30,531 (2.0%) of all nuclei come from rat, the human data drive the clustering. As expected the vast majority of the profiled nuclei (90.4%) came from muscle fiber (Fig. 2E).

We sought to independently assess cluster identity without relying on marker gene patterns and therefore focused on cluster-level TSS-distal ATAC-seq peaks, many of which would not be taken into account when assigning cell types using marker genes. We developed a logistic regression approach to score the similarity between these peaks and enhancer chromatin states from 127 Roadmap Epigenomics cell types (Fig. 2F) (31). We found concordance with the marker gene-based cell type assignment approach (Fig. 2G). Remarkably this approach worked relatively well in assigning rat nuclei, despite the fact that the number of nuclei per cluster for rat ranged between six and twenty for the smallest four cell types (Table S5; Fig. 2H).

The majority of the nuclei were assigned as type I or type II muscle fibers. Genes previously discovered to be preferentially expressed in type I vs. type II muscle fibers (13) were usually similarly preferentially expressed in our snRNA-seq data (Fig. S11), validating the quality of the data and accuracy of muscle fiber type assignments.

### Integration of cell-type-specific ATAC-seq peaks with UK Biobank GWAS reveals cell type roles in complex phenotypes

Genetic variants associated with complex traits and disease are frequently located in non-coding regions of the genome (33–35). Variants associated with a given complex trait are expected to be enriched specifically in non-coding regulatory elements of the trait-relevant cell types; for example, T2D-associated genetic variants are enriched in regulatory elements specific to pancreatic islets and beta cells (34,36–44), and variants associated with autoimmune disorders are enriched in immune cell-specific regulatory elements (36). Variant enrichment in cell-specific regulatory elements can therefore be used to determine which cell types are relevant to a given trait or disease. Variants in high linkage disequilibrium (LD) with trait-influencing SNPs are often statistically associated with the trait as well, making it difficult to infer the causal SNP through statistical association alone. Epigenomic data, such as chromatin accessibility in trait-relevant cell types, can be used to nominate causal genetic variants under the assumption that non-coding SNPs in accessible regions of the genome are more likely to be causally related to a trait than non-coding SNPs in inaccessible regions.

To explore the relationship between complex traits and the cell types present in our data, as well as demonstrate the value of our muscle cell type chromatin data in narrowing the post-GWAS search space, we used LD score regression (LDSC) (36,45) to perform a partitioned heritability analysis using GWAS of 404 heritable traits from the UK Biobank (46) (http://www.nealelab.is/uk-biobank/) and our muscle cell type open chromatin regions (Table S6; see Methods) (36,45). Results for all traits in which at least one of our cell types showed significant (P < 0.05) enrichment after Benjamini-Yekutieli correction are displayed in Fig. 3A. Due to the heavy multiple testing correction burden, relatively few traits meet this threshold. However, we observed immune cell abundance traits show enrichment for the immune cell cluster, and diastolic blood pressure GWAS SNPs are enriched in smooth muscle ATAC-seq peaks. In addition, we see that several skeletal trait GWAS SNPs are enriched in mesenchymal stem cell peaks. Previous work has shown a central role of bone mesenchymal stem cells in osteoblast development (47,48). In addition, SNPs for several corneal traits are also enriched in mesenchymal stem cell peaks, consistent with previously observed enrichment of corneal thickness GWAS SNPs in mesenchymal stem cell/connective tissue cell annotations (49). Results using rat peaks projected into human coordinates largely mirror the human mesenchymal stem cell enrichment findings (Fig. S12).

**Figure 3:**
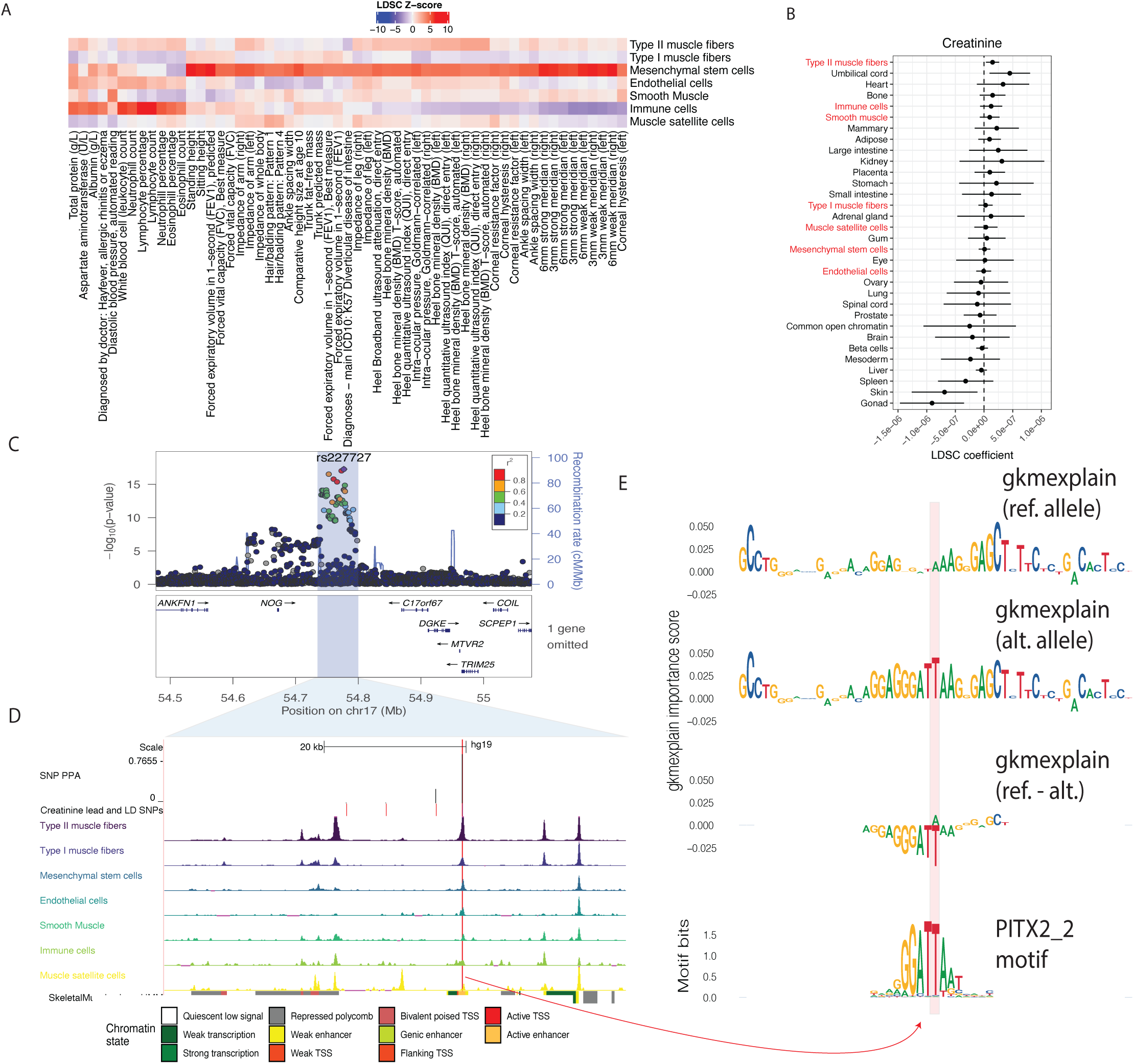
(A) UK Biobank LDSC partitioned heritability results for traits for which one of the muscle cell types was significant after Benjamini-Yekutieli correction. (B) LDSC partitioned heritability results for creatinine (UK Biobank trait 30700). Red y-axis labels refer to the muscle snATAC-seq cell type annotations. (C) Locuszoom plot for *C17orf67* locus in the UK Biobank creatinine GWAS. (D) ATAC-seq signal in the region highlighted in (C). The red line represents the location of SNP rs227727. All tracks are normalized to 1M reads. SNPs shown have LD > 0.8 with the lead SNP based on the European samples in 1000 Genomes Phase 3 (Version 5; 1000 Genomes Project Consortium et al., 2015). (E). gkmexplain importance scores for the ref and alt allele-containing sequences (top two rows), and the difference between the ref and alt allele importance scores (third row), which resembles the PITX2_2 motif predicted to be disrupted by the A allele (bottom row).

One muscle-related trait included in the UK Biobank is creatinine level. In humans most serum creatinine is produced by skeletal muscle and is filtered by the kidneys (50). Creatinine levels are commonly used as a biomarker for kidney function but correlate with muscle mass and have been used to score sarcopenia (51–53). In our enrichment analysis, the cell type with the highest LDSC coefficient Z-score was type II muscle fibers (z-score = 2.5; Fig. 3B).

Integrating the ATAC-seq results with the GWAS summary statistics can help nominate causal SNPs. One example is the *C17orf67* locus in the creatinine GWAS (Fig. 3C). The lead SNP at this locus (rs227727; p = 5.38e-18) lies in an intergenic region 92 kb from *C17orf67* and 104 kb from *NOG*. This SNP is in an ATAC-seq peak in several muscle cell types, though the signal is largest in type II muscle fibers (Fig. 3D). The peak corresponds to an enhancer chromatin state in muscle, amongst other cell types (31). We used the Probabilistic Identification of Causal SNPs (PICS) tool (54) to estimate the probability that nearby SNPs were causal given the pattern of linkage disequilibrium at the locus. PICS assigned the index SNP, rs227727, a probability of 0.766 of being the causal SNP. A tightly linked SNP, rs227731 (R^2^ = 0.99), had a probability of 0.221; no other SNPs had probability greater than 0.01. SNP rs227731 is not in an ATAC-seq peak in any of the muscle cell types we identified nor is it in any of ENCODE’s 1.3 million candidate cis-regulatory elements (55,56) or any of the approximately 3.6 million DNaseI hypersensitive sites (DHS) annotated in (57), suggesting that the index SNP rs227727 is indeed the causal SNP. A previous study found that the A allele of rs227727 was associated with higher activity in an allelic luciferase assay in both human fetal oral epithelial cells (GMSM-K) and murine osteoblastic cells (MC3T3) (58). To predict allelic effects at this SNP in type II muscle fibers, we trained a gapped-kmer support vector machine model (gkm-SVM) (59,60) to detect kmers associated with increased or decreased chromatin accessibility using the top ATAC-seq peaks for each of our cell types and then ran deltaSVM (61) to predict this SNP’s effect on chromatin accessibility. DeltaSVM predicts a SNP’s effect by comparing the gkm-SVM inferred kmer weights for kmers created by the reference vs the alt allele; we transformed the deltaSVM score to a z-score based on the distribution of the predicted impacts of all autosomal 1000 Genomes SNPs (62). The type II muscle fiber deltaSVM z-score for this SNP was 0.73 (directionally favoring the alt allele, T, having higher chromatin accessibility, although the z-score is not statistically significant). We also attempted to interpret how each allele of the SNP affects the gkm-SVM model’s score for the sequence using the gkmexplain software package, which scores the importance of each base in a sequence to the gkm-SVM model score for the sequence (63). We ran gkmexplain on the sequence surrounding the SNP in the presence of either the reference or the alternative allele and compared the results (Fig. 3E). The change in the gkmexplain importance scores in the presence of the reference vs alternative allele resembled several known homeodomain TF motifs predicted to be disrupted by the reference allele such as that of PITX2, suggesting that the alternate allele may have directionally (non-significant) greater predicted chromatin accessibility because it is a better match to these homeodomain TF motifs (Fig. 3E) (64). We note, however, that the deltaSVM z-score of the SNP as well as the gkmexplain importance scores of the SNP and surrounding nucleotides are of low magnitude, suggesting that the reference allele may reduce the binding of PITX2 or another homeodomain TF without such a dramatic effect on local chromatin accessibility. Biologically, the nearby *NOG* gene is a particularly compelling candidate target gene of this regulatory element, as its product (noggin) regulates BMP signaling and is involved in muscle growth and maintenance (65–70). Integrated with the GWAS summary statistics and these additional resources, our ATAC-seq data adds to existing evidence that SNP rs227727 alters the activity of a gene regulatory element and is a prime candidate to impact creatinine levels.

### Integration of cell type-specific ATAC-seq peaks with T2D GWAS credible sets nominates causal cell types, regulatory elements, and SNPs

It is well-established that T2D GWAS SNPs overlap pancreatic islet/beta cell enhancers (34,37,38,41,43); however, some SNPs may act through other T2D-relevant tissues, such as muscle, adipose, or liver. We therefore used LDSC to perform a partitioned heritability analysis for T2D-associated SNPs (38) in each of the muscle cell types as well as in beta cell ATAC-seq peaks, adipose ATAC-seq peaks, and liver DNaseI hypersensitive sites (see Methods) (Figs. 4A, S13A). When modeling each cell type separately (adjusting for the cell type-agnostic LDSC baseline annotations and common open chromatin regions), we found significant enrichment (after Bonferroni correction for 40 tests) in type II muscle fibers and beta cells, though when modeling all cell types in a single joint model only beta cell open chromatin regions showed significant enrichment (Fig. S13A). We performed a similar analysis on GWAS SNPs for a T2D-related trait, fasting insulin (Figs. 4A, S13A) (71). For fasting insulin, we found significant enrichment in mesenchymal stem cells, immune cells, and bulk adipose when modeling each cell type individually, but only adipose showed significant enrichment when modeling all cell types jointly. For fasting insulin, we note that the small sample size of that GWAS means the analysis was likely underpowered, leaving open the possibility that other cell types will show significant enrichment when GWAS with larger sample sizes are available. We also note that the adipose open chromatin regions are derived from bulk tissue open chromatin profiling; it is therefore possible that at least some of the signal from adipose is being driven by cell types shared between our muscle samples and adipose tissue, such as mesenchymal stem cells. This is an area for further exploration when single-cell/single-nucleus data from adipose is available.

**Figure 4:**
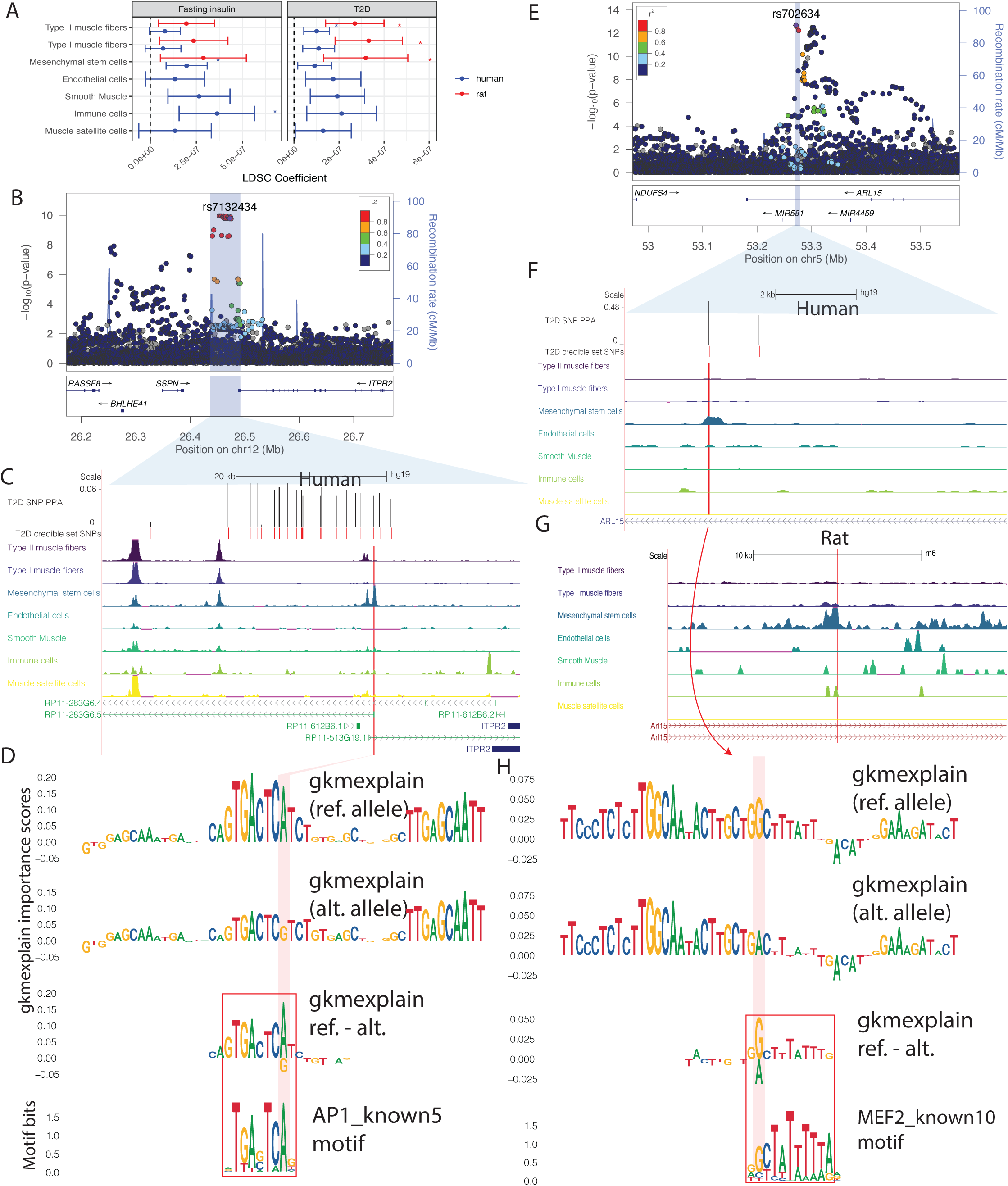
(A) LDSC partitioned heritability results for T2D (BMI-unadjusted) and Fasting insulin GWAS (BMI-adjusted), using human peak calls. For each of the cell types, one model was run adjusting for cell type-agnostic annotations from the LDSC baseline model and common open chromatin regions. Asterisks represent Bonferroni significance (p < 0.05 after adjusting for 40 tests). (B) locuszoom plot for *ITPR2* locus in the DIAMANTE data. (C) DIAMANTE credible set near the *ITPR2* gene, consisting of 22 SNPs. One SNP (rs7132434; highlighted in red) overlaps a peak call in any of the muscle cell types. (D) gkmexplain importance scores for the ref and alt allele (top two rows) and the difference between the ref and alt importance scores (third row); the G allele disrupts an AP1 motif (bottom row). (E). locuszoom plot for *ARL15* locus in the DIAMANTE data. (F). DIAMANTE credible set SNPs near the *ARL15* gene. The three SNPs represent the three-SNP credible set discussed in the text. One of these SNPs (rs702634; highlighted in red) overlaps a mesenchymal stem cell specific peak. (G). Projecting the SNP highlighted in (F), rs702634, into the rat genome (projected SNP position indicated by the red vertical line) shows the corresponding region has open chromatin in rat mesenchymal stem cells. (H). gkmexplain importance scores for the ref and alt alleles (top two rows), the difference between them (third row), and a MEF2 motif disrupted by rs702634.

We performed similar GWAS enrichments using the rat muscle cell type peaks projected into human coordinates (Fig. 4A, S13B). For T2D we found muscle fiber types and mesenchymal stem cells were significantly enriched after Bonferroni correction, but as with human muscle cell types these enrichments did not persist in a joint model with all cell types (Fig S13B). For fasting insulin no rat muscle cell types showed enrichment after Bonferroni correction.

While none of our cell types showed significant enrichment in 10-cell-type models after Bonferroni correction, it is still possible that some T2D GWAS loci act through muscle cell types or cell types shared between muscle and other tissues such as adipose. There are a substantial number of T2D GWAS credible sets that show no overlap with pancreatic islet functional annotations (38). We therefore overlapped 380 previously-published T2D GWAS signals with 99% genetic credible set SNPs (38) with our snATAC-seq peaks to nominate SNPs that may be acting through the muscle cell types, including those that are expected to be shared with adipose (Table S7).

One locus highlighted by our data is the *ITPR2* locus on chromosome 12 (Fig. 4B). This locus contains 22 credible set SNPs, none with a particularly high posterior probability of association (PPA) in the DIAMANTE genetic fine-mapping (maximum across all credible set SNPs = 0.06). Only one SNP (rs7132434; PPA = 0.042) overlaps any of our muscle cell type peak calls (Fig. 4C). This SNP is in a large mesenchymal stem cell ATAC-seq peak, and also overlaps peak calls in smooth muscle and blood, though the chromatin accessibility signal in those cell types is lower in our data. The SNP also overlaps a peak call in a subset of adipose and islet samples (Fig. S14). We found that this SNP had a large deltaSVM z-score in several of the muscle cell types (absolute z-score = 2.88 in mesenchymal stem cells; the T2D risk allele, A, is predicted to result in greater chromatin accessibility). We ran gkmexplain on the sequence surrounding the SNP and found the gkmexplain importance scores for the sequence in the presence of the risk allele resembled an AP-1 motif (Fig. 4D) (64). A literature search revealed that the element underlying this SNP has been validated for enhancer activity using a luciferase assay (in the 786-O cell line) and the risk allele showed preferential binding of the AP-1 transcription factor in an EMSA assay in the same study and cell line (Bigot et al., 2016), consistent with our findings. We note that this SNP is also a 95% credible set SNP for waist-hip ratio (one of eight SNPs in the credible set) (72). We therefore hypothesize that rs7132434 is the causal SNP at this locus, and that it may be acting through mesenchymal stem cells.

A second locus highlighted by our data is an intronic locus in the *ARL15* gene (Fig 4E). The DIAMANTE genetic fine-mapping narrowed the list of potentially causal SNPs at this locus to three (two other, larger DIAMANTE genetic fine-mapping credible sets are also annotated to *ARL15*). SNPs in this credible set are statistically associated with fasting insulin (73), and more broadly variants in or near *ARL15* associate with metabolic traits including adiponectin, HDL cholesterol levels, and BMI (73–75), suggesting that the locus may affect T2D risk not through islets but through adipose or a related cell type. Interestingly, none of the SNPs overlap with any of ENCODE’s 1.3 million candidate cis-regulatory elements (55,56) or any of the approximately 3.6 million DNaseI hypersensitive sites (DHS) annotated in (57); however, in our data we find that one of the SNPs (rs702634) is in the center of a mesenchymal stem cell specific ATAC-seq peak (Fig. 4F), and a mesenchymal stem cell peak is likewise present in the corresponding position in the rat genome (Fig. 4G), indicating that this is a regulatory element that has been conserved across species. The DIAMANTE genetic fine-mapping assigned this SNP a probability of 0.48 of being the causal SNP at this locus, higher than either of the other two SNPs (0.33 and 0.19, respectively). We examined publicly-available beta cell (n = 1), islet (n = 10) (41), and adipose (n = 3) (76) ATAC-seq data to see if hints of this peak are present in these T2D-relevant cell types. No convincing signal appears to be present in beta cell or islet data; a weak increase in signal at that SNP is evident in the adipose samples and a peak is called (Fig. S15). As mesenchymal stem cells are one component of adipose tissue, it is possible that the weak signal in adipose is due to mesenchymal stem cell populations within adipose; this is one area for follow-up when adipose single-nucleus ATAC-seq data is available. The absolute deltaSVM z-score in mesenchymal stem cells for this SNP was 0.48, indicating it does not have a large impact on predicted chromatin accessibility; however, the risk allele is predicted to disrupt a MEF2 motif (64,77), and we found the change in gkmexplain importance scores between the reference and alternative allele showed similarity to this motif (Fig. 4H). This data is consistent with a model in which rs702634 is the causal SNP and acts through mesenchymal stem cells.

## Discussion

Here we present snATAC-seq and snRNA-seq for human skeletal muscle and snATAC-seq for rat skeletal muscle, which we use to map the transcriptomes and chromatin accessibility of cell types present in skeletal muscle samples. The cell types identified are consistent with known biology and with previous studies of human (13) and mouse (16,17,20) skeletal muscle tissue. However, our use of single-nucleus rather than single-cell techniques allows us to capture muscle fiber nuclei, cell types missing from previously published snRNA-seq datasets. To our knowledge this is the first published snATAC-seq dataset for human and rat skeletal muscle tissue. We therefore anticipate that this dataset will be useful in nominating causal GWAS SNPs and demonstrate this by integrating the data with UK Biobank and previously published T2D GWAS credible sets, highlighting potentially causal SNPs at the *NOG, ARL15*, and *ITPR2* loci.

Additionally, we explore the effect of two technical parameters on snRNA-seq and snATAC-seq results. First, we find that FANS (using DRAQ7 staining) substantially alters snATAC-seq results. Though the stereotypical ATAC-seq fragment length distribution is observed, signal-to-noise (as measured by TSS enrichment and fraction of reads in peaks, as well as by visual inspection) appears to decrease substantially relative to non-FANS libraries. We note that the effect of FANS (nucleus sorting) may differ from that of FACS (cell sorting). snRNA-seq results appear to be substantially less sensitive to FANS -- the pseudobulk gene expression from FANS libraries correlates strongly with that from non-FANS libraries -- suggesting that chromatin is more sensitive to FANS than is RNA. We also observed higher nucleus yield in our FANS snRNA-seq libraries than our non-FANS libraries. There are several potential explanations for this. One is that the nuclei counting step that necessarily precedes loading of the 10X platform may be sensitive to debris. If greater amounts of debris are observed in non-FANS libraries, nucleus concentration may be systematically overestimated in non-FANS libraries, resulting in more nuclei actually being loaded onto the 10X platform from FANS libraries. While not mutually exclusive, FANS may also decrease the amount of debris being loaded into the 10X platform, and thereby improve nucleus capture.

We found snATAC-seq and snRNA-seq results were remarkably consistent at different loading concentrations. One clear caveat is that this may change as the loading concentration is further reduced or increased. It is also important to note that the actual number of nuclei loaded may differ from the estimated 20k or 40k nuclei. As discussed above, it is possible that debris in the input preparation makes nucleus counting less accurate, in which case our cited values may not reflect the true values. However, because the same nuclear preparation was used as input for the 20k and 40k nuclei libraries, the two-fold difference in loading concentration should be reliable, even if the absolute values are skewed.

The GWAS enrichments presented here will be one interesting area to follow up on as more snATAC-seq data is published. Interpretation of the results is complicated by the fact that many tissues share cell types. For example, mesenchymal stem cell-like populations exist in many tissues besides muscle, such as adipose tissue and bone marrow. Taking the fasting insulin enrichments as an example, we found that the enrichment of GWAS SNPs in muscle cell type ATAC-seq peaks disappeared when adipose tissue was included in the enrichment model. However, it is possible that the adipose enrichment is being driven in part by mesenchymal stem cell populations within adipose itself. Direct comparison of snATAC-seq and snRNA-seq profiles from mesenchymal stem cells from a wider array of tissues will help tease apart commonalities and tissue-specific differences in this interesting population.

## Methods

### Reproducibility of computational analyses

Code used for analyses in this manuscript are available at https://github.com/ParkerLab/2020-sn-muscle.

### snATAC-seq and snRNA-seq, FANS vs no FANS experiment

Three separate pieces of tissue were cut from a single human skeletal muscle sample (weighing 60mg, 50mg and 50mg; sample HSM1, quadriceps femoris muscle group). Nuclei were isolated using a modified version of the ENCODE protocol (protocol S1) (56,56), customized from Step 5 onwards to accommodate FANS (Fluorescence assisted nuclei sorting). In step 5, the nuclei were resuspended in 700 µL of Sort buffer (1% BSA, 1mM EDTA in PBS) and filtered through a 30 µm filter. Three different nuclei isolations were performed and the nuclei suspended in sort buffer were mixed, pooled together and divided into two groups, one with FANS and one without FANS. FANS nuclei were sorted according to the previously published FANS protocol using DRAQ7 (78). DRAQ7 (0.3mM from Cell Signaling Technology) was added to the FANS nuclei suspension, at 100 fold dilution to get a final concentration of 3 μM. Nuclei were gently mixed and incubated for 10 minutes on ice. Nuclei were analyzed in the presence of DRAQ7 and sorted for high DRAQ7 positive signal using Beckman Coulter’s Astrios MoFlo. We followed the gating strategy outlined in the FANS protocol (Preissl et al, 2018). The sorted nuclei were collected in a recovery buffer (5% BSA in PBS). The nuclei with and without FANS were spun at 1000g for 15 min at 4°C. The nuclei were resuspended in 100 µL of 1X diluted nuclei buffer and counted in the Countess II FL Automated Cell Counter. The appropriate amount of nuclei were split for snRNA-seq and spun down at 500g for 10 min at 4°C and resuspended in RNA nuclei buffer (1%BSA+PBS in 0.2U RNAse inhibitor). The nuclei at appropriate concentration for snATAC-seq and snRNA-seq were submitted to the Advanced Genomics core for all the snATAC-seq and snRNA-seq processing on the 10X Genomics Chromium platform (v. 3.1 chemistry for snRNA-seq). For each modality nuclei were loaded at 15.4K nuclei/well.

### snATAC-seq and snRNA-seq, loading 20k or 40k nuclei

Two pieces of tissue (weighing 85.3 mg and 85.8 mg) were cut from one human skeletal muscle sample (HSM1) and two tissue pieces (weighing 95.9 mg and 92.6 mg) were cut from a second human skeletal muscle sample (HSM2; quadriceps femoris muscle group). Each of the samples was cut on dry ice using a frozen scalpel to prevent thawing. The samples were pulverized using a CP02 cryoPREP automated dry pulverizer (Covaris 500001). We developed a customized protocol (protocol S2) derived from the previously published ENCODE protocol (56,56) and used it to isolate nuclei, which is compatible with both snATAC-seq and snRNA-seq. All four pulverized tissues pieces were mixed and redistributed to perform four different nuclei isolations. The desired concentration of nuclei was achieved by resuspending the appropriate number of nuclei in 1X diluted nuclei buffer for snATAC-seq and RNA nuclei buffer (1% BSA in PBS containing 0.2U/uL of RNAse inhibitor) for snRNA-seq. The nuclei at appropriate concentration for snATAC-seq and snRNA-seq were submitted to the Advanced Genomics core for all the snATAC-seq and snRNA-seq processing on the 10X Genomics Chromium platform (v. 3.1 chemistry for snRNA-seq). For each modality nuclei were loaded at two different concentrations, 20K nuclei/well and 40K nuclei/well.

### snATAC-seq, human and rat mixed library

Tissue from human (49mg of pulverized human skeletal muscle; sample HSM1) and rat (45mg of pulverized gastrocnemius samples) were used in this single nuclei ATAC experiment. We used the previously published ENCODE protocol (protocol S1) (56,56) to isolate nuclei, which is compatible with both snATAC-seq and snRNA-seq. After isolating nuclei from each sample (species) individually, the nuclei were mixed in equal proportions. The desired concentration of nuclei was achieved by resuspending the appropriate number of nuclei in 1X diluted nuclei buffer for snATAC-seq. The nuclei at the appropriate concentration for snATAC were submitted to the Advanced Genomics core for all the snATAC-seq processing on the 10X Genomics Chromium platform. 15.4K nuclei were loaded into a single well.

### Bulk ATAC-seq

2 tissue pieces weighing 99.4 mg and 80.7 mg were cut from one human skeletal muscle sample (HSM1) and 2 pieces weighing 67.6 mg and 103.5 mg were cut from a second human skeletal muscle sample (HSM2). Each of the samples was cut on dry ice using frozen scalpel to prevent thawing. The samples were pulverized using a CP02 cryoPREP automated dry pulverizer (Covaris 500001). For bulk ATAC seq we followed the nuclei isolation protocol outlined in protocol S2, except in the final step the nuclei were resuspended in 250 μL of 1% BSA. The nuclei were counted in Countess II FL Automated Cell Counter, and the appropriate volume of the suspension for 50K nuclei was spun down and used for the downstream transposition reaction (a modified version of the ENCODE protocol; protocol S3) (56,56).

### Processing of muscle bulk ATAC-seq data

Adapters were trimmed using cta (v. 0.1.2; https://github.com/ParkerLab/cta). Reads were mapped to hg19 using bwa mem (-I 200,200,5000 -M; v. 0.7.15-r1140) (79). Duplicates were marked using picard MarkDuplicates (v. 2.21.3; https://broadinstitute.github.io/picard/). We used samtools to filter to high-quality, properly-paired autosomal read pairs (-f 3 -F 4 -F 8 -F 256 -F 1024 -F 2048 -q 30; v. 1.9 using htslib v. 1.9) (80). To call peaks, we used bedtools bamtobed to convert to a bed file (v. 2.27.1) and then used that file as input to MACS2 callpeak (--nomodel -- shift -100 --seed 762873 --extsize 200 --broad --keep-dup all --SPMR; v. 2.1.1.20160309) (81,82). To visualize the signal, we converted the bedgraph files output by MACS2 to bigwig files using bedGraphToBigWig (v. 4) (83).

### Processing of snATAC-seq data

Adapters were trimmed using cta. We used a custom python script (available in the GitHub repo) for barcode correction. Barcodes were corrected in a similar manner as in the 10X Genomics Cell Ranger ATAC v. 1.0 software. In brief, barcodes were checked against the 10X Genomics whitelist. If a barcode was not on the whitelist, then we found all whitelisted barcodes within a hamming distance of two from the bad barcode. For each of these whitelisted barcodes, we calculated the probability that the bad barcode should be assigned to the whitelisted barcode using the phred scores of the mismatched base(s) and the prior probability of a read coming from the whitelisted barcode (based on the whitelisted barcode’s abundance in the rest of the data). If there was at least a 97.5% chance that the bad barcode was derived from one specific whitelisted barcode, it was corrected to the whitelisted barcode.

Reads were mapped using bwa mem with flags ‘-I 200,200,5000 -M’. We used Picard MarkDuplicates to mark duplicates, and filtered to high-quality, non-duplicate autosomal read pairs using samtools view with flags ‘-f 3 -F 4 -F 8 -F 256 -F 1024 -F 2048 -q 30’. Quality control metrics were gathered on a per-nucleus basis using ataqv (v. 1.1.1) on the bam file with duplicates marked. In the case of the mixed rat and human snATAC-seq library, all reads were mapped to the hg19 and rn6 genomes separately, and then a nucleus was assigned as either rat or human by counting the number of high-quality, non-duplicate autosomal reads after mapping to either genome. If at least three times as many high-quality reads were present after mapping to one genome than to the other, the nucleus was assigned to either the rat or human sample as appropriate. In the case that fewer than three times as many high-quality reads mapped to one genome as to the other, the nucleus was not assigned to either species and was dropped.

For the two snATAC-seq libraries that contained a mix of nuclei from the two human individuals, we assigned nuclei to biological samples (and determined doublets) using demuxlet (84) with SNP calls from the bulk ATAC-seq libraries. To call SNPs on the bulk ATAC-seq bam files, we first merged the two bulk technical replicate ATAC-seq bam files for each individual, then filtered out reads with edit distance > 2 from the hg19 reference. Used samtools mpileup (-R -Q 20 -d 10000 -E) on these two bam files as input to bcftools call (-v -f GQ; v. 1.9). We then used bcftools filter to filter to those positions where both individuals had genotype quality (GQ) > 90. This VCF file was used as input to demuxlet (option ‘--field PL’; git commit b7453fc, modified as described in GitHub issue #15).

When comparing aggregate snATAC-seq signal to bulk ATAC-seq signal (Fig. 1), we eliminated sequencing reads corresponding to nucleus barcodes that couldn’t be matched to the 10X barcode whitelist, but otherwise processed it as bulk ATAC-seq data (i.e., marking duplicates ignoring cell-level information, and not filtering to quality nuclei).

To select quality nuclei from each library, we selected nuclei (barcodes) meeting the thresholds in Table S1. In addition to setting a threshold for minimum fragments (to filter out barcodes that only capture ambient DNA fragments), we set a threshold for maximum fragments, because barcodes with very high fragment counts may be enriched for doublets (41). We also set a threshold for minimum TSS enrichment (because ATAC-seq signal for healthy nuclei is expected to be enriched near TSS (41,85,86)), and we filtered out barcodes that showed an unexpectedly large fraction of reads coming from a single autosome (see (29)).

### Processing of snRNA-seq data

snRNA-seq data was processed using starSOLO (STAR v. 2.7.3a), which outputs the count matrices needed for most of the analyses (87). To select quality nuclei from each library, we selected nuclei meeting the thresholds in Table S2 (we set a threshold for minimum UMIs to filter out barcodes that only capture ambient RNA; a threshold for maximum fragments, since barcodes with very high UMI counts may be enriched for doublets; and a threshold for maximum mitochondrial contamination, since barcodes with quality nuclei and low ambient RNA should show reduced mitochondrial contamination (88)). We used souporcell (as contained in the Singularity container downloaded from the souporcell GitHub on Dec. 10, 2019, and setting -k 2) to detect doublets in the libraries that were a mix of nuclei from two human individuals (89). We additionally ran doubletfinder (v. 2.0.2) (32) on each of the snRNA-seq libraries, and removed any nuclei that were called as a doublet by either souporcell or doubletfinder. When running Seurat (v. 3.0.2) for doubletfinder, we set selection.method = “vst” and nfeatures = 2000, and used the top 20 PCs to find neighbors and resolution = 0.8 to find clusters (90,91). When calling the doubletFinder_v3 function, we selected the doubletfinder pK based on the maximum ‘BCmetric’ after running the paramSweep_v3 function, set nExp assuming a 7.5% doublet rate (adjusting for the homotypic proportion as in the doubletfinder documentation example), and used the top 20 PCs.

### Clustering with LIGER

Nuclei were clustered using LIGER (v. 0.4.2; with R v. 3.5.1 and Seurat v. 2.3.0) (90–92). For snATAC-seq libraries, per-gene scores were computed by calculating the number of reads overlapping with each gene’s promoter/gene body using bedtools intersect. Gene promoter/body were calculated based on NCBI annotation GTF files (NCBI Rattus norvegicus Annotation Release 106 and Homo sapiens Updated Annotation Release 105.20190906), filtered to include only protein-coding/lncRNA genes with source ‘BestRefSeq’/BestRefSeq%2CGnomon’/’Curated Genomic’. Genes assigned to multiple chromosomes/strands were excluded, and then the regions for each gene were merged to get the gene body. Promoters were taken as the 3kb upstream of the TSS; after this, genes represented by multiple non-contiguous genomic stretches were excluded. For input to LIGER, all count matrices for a given modality and biological sample were concatenated together, so that there was 1 rat snATAC matrix, 2 human snATAC matrices, and 2 human snRNA matrices. For factorization, we used k = 15, lambda = 5, and nrep=5, using the smaller human snRNA matrix to select variable genes (as all the nuclei for that matrix were processed on a single day, and should therefore reflect less technical variation). For each of the downstream steps we dropped factors 3 and 5, as these had highly-loading ribosomal genes or showed relatively high specificity for one of the two omics modalities. For normalization, we set knnk (and small.clust.thresh) to 10 and resolution to 0.05, and centered the data. For the UMAP, we used n_neighbors = 15. We then called the clusterLouvainJaccard function to re-cluster cells using the normalized factors, with k = 17, and resolution = 0.05.

### Per-cluster processing of snATAC-seq data

The filtered reads from all snATAC-seq nuclei in each cluster were merged using samtools merge. Peaks were called and bigwig files produced as described for the bulk ATAC-seq data. Peak files were filtered against blacklist files available from http://hgdownload.cse.ucsc.edu/goldenPath/hg19/encodeDCC/wgEncodeMapability/wgEncode DacMapabilityConsensusExcludable.bed.gz and http://hgdownload.cse.ucsc.edu/goldenPath/hg19/encodeDCC/wgEncodeMapability/wgEncode DukeMapabilityRegionsExcludable.bed.gz (hg19) (56) and https://github.com/shwetaramdas/maskfiles/tree/master/rataccessibleregionsmaskfiles/strains_intersect.bed for rn6 (93).

For analysis of rat peak overlap with human GWAS data, rat peaks were projected into the human genome using bnMapper (v. 0.8.6) and the chain file at http://hgdownload.cse.ucsc.edu/goldenpath/rn6/liftOver/rn6ToHg19.over.chain.gz.

### Roadmap enhancer regression

We called peaks on the aggregate of the nuclei in each cluster, and then took the union of peaks across all clusters to generate a master peak list. We then used logistic regression to model, for each cluster and each Roadmap Epigenomics cell type in the Roadmap 15-state chromHMM model, the accessibility of each TSS-distal master peak (> 5kb from a RefSeq TSS) in that cluster as a function of the posterior probability that that master peak is an enhancer in that Roadmap cell type according to the Roadmap chromHMM model (31). Since the posteriors are given in 200 bp windows, and there are also 3 different enhancer states (‘Genic enhancers’, ‘Enhancers’, and ‘Bivalent Enhancer’), multiple windows overlap with each master peak -- the posterior for the master peak is therefore taken as the maximum of the 200 bp window posteriors, across all 3 of the enhancer states. The model coefficient was used as the (unnormalized) score for that Roadmap cell type in that cluster, and the normalized score was simply the score for that Roadmap cell type in that cluster divided by the max score across all cell types for that cluster. For rat peaks, in addition to removing master peaks near TSS in rat coordinates, we additionally removed master peaks that were within 5 kb of a TSS after projecting into human coordinates.

### Non-muscle cell type open chromatin annotations used in GWAS

To create the adipose open chromatin regions, we processed the three adipose ATAC-seq libraries from (76). Adapter sequences were removed using Cutadapt (v. 1.12) (94) before mapping to hg19 with bwa mem (-I 200,200,5000 -M). Duplicates were marked using picard MarkDuplicates and BAM files were filtered using samtools view (-F 4 -F 256 -F 1024 -F 2048 -q 30) before converting to BED format (bamtools bamtobed) and calling peaks with MACS2 (-- nomodel --shift -100 --seed 2018 --extsize 200 --broad --keep-dup all --SPMR). We then took the union of peaks across the three samples, keeping those merged peaks that appeared in at least two samples.

The beta cell ATAC-seq peaks were taken from (41). We used the peaks called using all beta cell nuclei.

Common open chromatin regions were derived from the DNaseI hypersensitive sites from (57). The DHS index from Meuleman et al. was downloaded from https://www.meuleman.org/DHS_Index_and_Vocabulary_hg38_WM20190703.txt.gz on March 21, 2020. We lifted open chromatin regions from hg38 to hg19 using liftOver with the chain file from http://hgdownload.cse.ucsc.edu/goldenpath/hg38/liftOver/hg38ToHg19.over.chain.gz (95). We then kept those that were labeled as ‘tissue invariant’ and that appeared in at least 500 of the 733 samples.

We also used open chromatin regions from (57) for adrenal gland, bone, brain, eye, gonad, gum, heart, kidney, large intestine, liver, lung, mammary, mesoderm, ovary, placenta, prostate, skin, small intestine, spinal cord, spleen, stomach, and umbilical cord. For each tissue, we took the non-cancerous samples labeled ‘Primary’ from that tissue and kept those DNaseI hypersensitive sites that appeared in at least 50% of the samples from that tissue.

### UK Biobank GWAS enrichment

We downloaded UK Biobank GWAS summary statistics made available by the Benjamin Neale lab (v2 of their analysis, initially made public on August 1, 2018; http://www.nealelab.is/uk-biobank/) (46). Specifically, we downloaded the ‘both sex’ GWAS summary statistic files listed in the ‘UKBB GWAS Imputed v3 - File Manifest Release 20180731’ spreadsheet available at https://docs.google.com/spreadsheets/d/1kvPoupSzsSFBNSztMzl04xMoSC3Kcx3CrjVf4yBmES U (downloaded on April 9, 2020). Because some traits may not be appropriate for such an enrichment analysis (because they are not strongly polygenic, because the phenotypes are untrustworthy, etc.), we kept only traits deemed as ‘high confidence’ and with estimated heritability > 0.01 (and z-score > 7) based on the Neale Lab’s own LD score regression heritability analysis of the GWAS results. Their rating criteria are described on their UKBB LDSC GitHub page (https://nealelab.github.io/UKBB_ldsc/confidence.html) and their LD score regression results (with confidence ratings) were downloaded from https://www.dropbox.com/s/ipeqyhrpdqav5uh/ukb31063_h2_all.02Oct2019.tsv.gz?dl=1. For each trait, we used the ‘primary’ GWAS result, as indicated in that file. Any traits that did not have a combined male and female GWAS analysis were dropped. The creatinine GWAS highlighted in the text was trait 30700_irnt (“Creatinine (quantile)”).

The LDSC software package (v. 1.0.1) includes a ‘baseline’ model with 59 categories derived from 28 genomic annotations (36,45). Many of these annotations are cell type agnostic; e.g. a SNP’s minor allele frequency does not change between cell types. However, other annotations in the baseline model are not cell type agnostic; for example, the FANTOM5 enhancer annotation is derived from experiments performed on a range of different cell types, and may change substantially if the cell types used to create the annotation were to change. When performing the UK Biobank GWAS enrichments, we utilized the cell-type agnostic annotations from the LDCS baseline model (Table S8). In order to reduce the likelihood of model misspecification, we then added common open chromatin regions and open chromatin regions from a range of cell types. Specifically, we added (1) beta cell ATAC-seq peaks, (2) adipose ATAC-seq peaks, (3) DNase-seq peaks derived from the 22 tissues/organs listed above, and (4) the ATAC-seq peaks from all seven of our snATAC-seq cell types. The various annotation files (regression weights, frequencies, etc.) required for running LDSC were downloaded from https://data.broadinstitute.org/alkesgroup/LDSCORE. LD scores were calculated using the Phase 3 1000 Genomes data, keeping only the HapMap3 SNPs as recommended by the LDSC authors and using only SNPs with minimum MAF of 0.01. GWAS summary statistics were prepared for LDSC using the munge_sumstats.py script, with option --merge-alleles w_hm3.snplist (where w_hm3.snplist is the file in the data download). When running the regression, we required a minimum MAF of 0.05, and utilized the Phase 3 1000 Genomes SNP frequencies/weights.

### T2D and fasting insulin GWAS enrichment

We used the T2D (BMI unadjusted) and fasting insulin (BMI adjusted) GWAS summary statistics from (Mahajan et al., 2018) and (Manning et al., 2012), respectively.

Because the cell types relevant to T2D are generally thought to be pancreatic beta cells, adipose, muscle, and liver, we performed enrichments using each of these cell types, common open chromatin, and the cell type-agnostic LDSC baseline annotations. First, for each of these muscle/beta cell/adipose/liver cell types, we ran one model containing the open chromatin from that cell type, the common open chromatin regions, and the cell type-agnostic LDSC baseline annotations. Then, we ran one joint model containing all of those cell types and annotations. LDSC parameters were the same as for the UK Biobank GWAS enrichments.

### T2D GWAS locus genome browser screenshots and peak overlaps

All signal tracks in the genome browser were created by converting the normalized bedgraph files output by MACS2 to bigwig files using bedGraphToBigWig (v. 4) (83).

Processing and provenance of adipose ATAC-seq and beta cell ATAC-seq is described above. The 10 bulk islet libraries were from (41). These libraries were processed as described in that manuscript, except we used the 10% FDR peak set from peak calling on the unsubsampled libraries.

### Predicting SNP regulatory impact

We used the lsgkm package modified by the Kundaje lab with gkmexplain (https://github.com/kundajelab/lsgkm; commit c3758d5bee7) (59,60,63). For each cell type, we took the 150 bps on either side of the summits of the top 40,000 narrowPeaks (by p-value) as the positive sequences for gkmSVM. To generate negative sequences, we took windows across the genome (step size = 200), removed those containing Ns, overlapping hg19 blacklists, overlapping any FDR 10% broadPeaks from that cell type, or having repeat content > 60%, and then for each positive sequence selected a negative sequence with matching GC content and repeat content (repeat content was calculated based on the hg19 simpleRepeat table from the UCSC genome browser (96,97), downloaded on March 29, 2020, which contains simple tandem repeats annotated by Tandem Repeats Finder (98); GC content and repeat content for the negative sequence was required to be within 2% of that of the positive sequence; in the case that no such negative sequence could be found, the positive sequence was dropped from the analysis). We held out 15% of sequences as test data, and trained the gkmSVM model on the remaining 85% of sequences, setting l = 10 and k = 6 and using the gkm kernel. Using this model and deltaSVM (61), we predicted the effect of all autosomal 1000 Genomes phase 3 SNPs (downloaded on May 27, 2015 from ftp://ftp.1000genomes.ebi.ac.uk/vol1/ftp/release/20130502) (62). For each muscle cell type, deltaSVM scores were converted to z-scores based on the distribution of scores across all SNPs for that cell type. We additionally passed the gkmSVM model to gkmexplain to generate importance scores for sequences containing the ref/alt alleles.

### Overlap of SNPs and peaks with ENCODE candidate cis-regulatory elements

The set of 1,310,152 candidate cis-regulatory elements in ENCODE’s ‘Registry of candidate Regulatory Elements’ (in hg19 coordinates) were fetched from the ENCODE web portal on April 7, 2020 (55,56).

### Locuszoom plots

Locuszoom plots were created for the DIAMANTE T2D GWAS summary statistics with the locuszoom standalone v. 1.4, using the Nov. 2014 EUR 1000 Genomes data included in the download (--pop EUR --source 1000G_Nov2014) (99).

### PICS

We used the online PICS tool (54) (https://pubs.broadinstitute.org/pubs/finemapping/pics.php) with the EUR LD structure. The tool was accessed on April 13, 2020.

### Motif scan

The MEF2 motif scan was performed using FIMO (v. 5.0.4) with a background model calculated from the hg19 reference genome (77).

## Supporting information

Protocol S1

Protocol S2

Protocol S3

## Declarations

### Ethics approval and consent to participate

Human samples were approved by the University of Michigan IRB protocol # HUM 000060733. Collection of the rat muscle sample was approved by the University of Michigan Institutional Animal Care and Use Committee.

### Consent for publication

Not applicable.

### Availability of data and materials

Raw sequencing reads generated during this study are not publicly available due to privacy restrictions. Processed data is available in a Zenodo repository (10.5281/zenodo.3926660).

### Authors’ contributions

NM generated the bulk ATAC-seq data and performed the nuclear isolations for the single-nucleus datasets. AV and VR processed existing islet and adipose ATAC-seq data. JK helped set up gkm-SVM models. CL helped coordinate production of the single-nucleus data. CFB provided the rat muscle sample, and KG provided human muscle samples. PO performed all computational processing and analyses of the data not attributed to others, and contributed to manuscript writing. SCJP designed and supervised the study and contributed to manuscript writing. All authors read and approved the manuscript.

## Acknowledgements

We wish to thank the University of Michigan Advanced Genomics Core for their assistance in generating the snRNA-seq and snATAC-seq libraries, and the University of Michigan Flow Cytometry Core for their help performing FANS. We are grateful to the Benjamin Neale lab for providing their UK Biobank GWAS and LDSC results to the scientific community. We also thank members of the Parker lab for their helpful feedback.

## Figure legends

**Figure S1:**
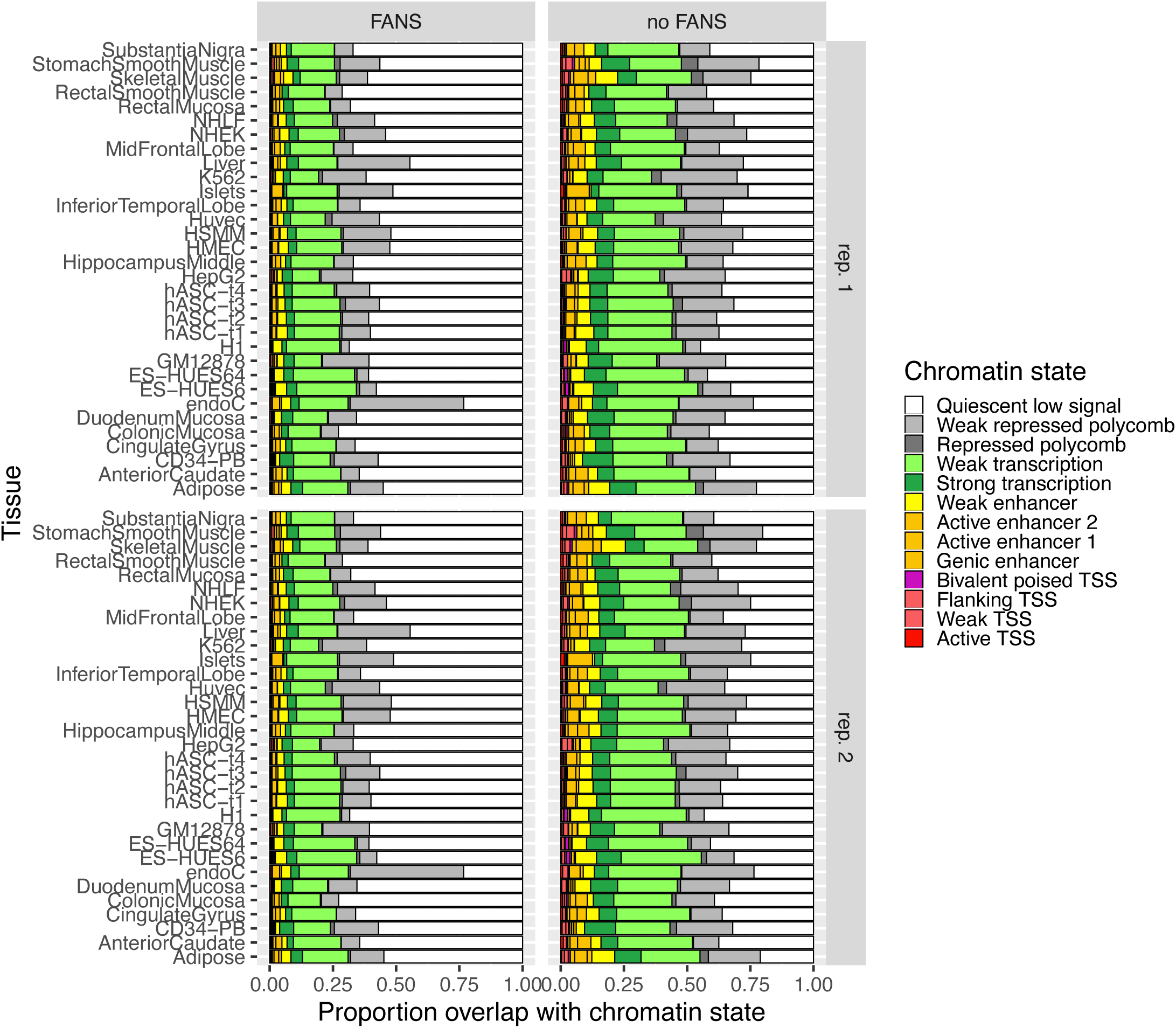
Chromatin state overlap for TSS-distal (> 5kb from TSS) ATAC-seq peaks from the FANS and non-FANS snATAC-seq libraries.

**Figure S2:**
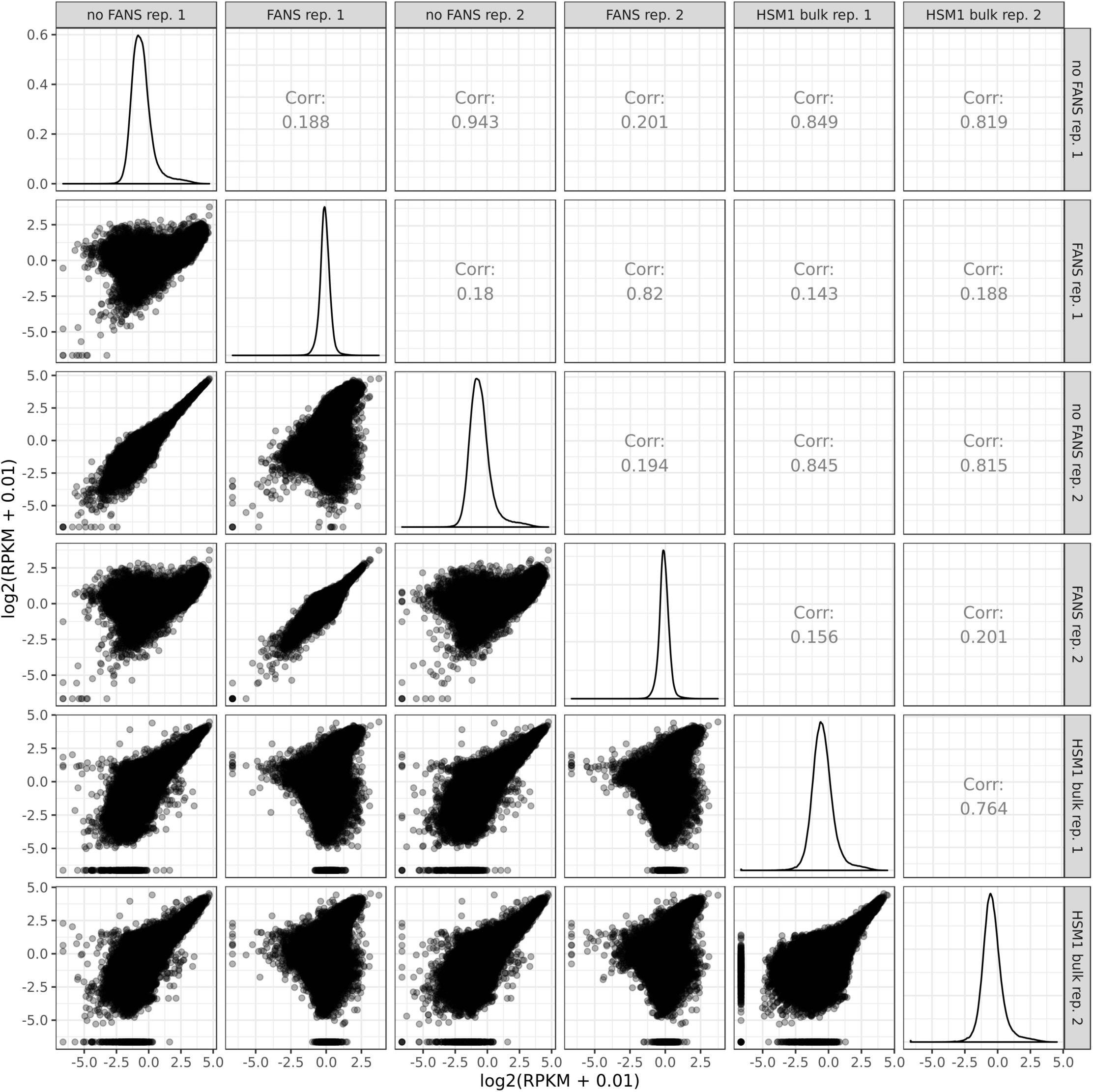
Correlation between FANS snATAC-seq, non-FANS snATAC-seq, and standard bulk ATAC-seq libraries. Each point represents one peak.

**Figure S3:**
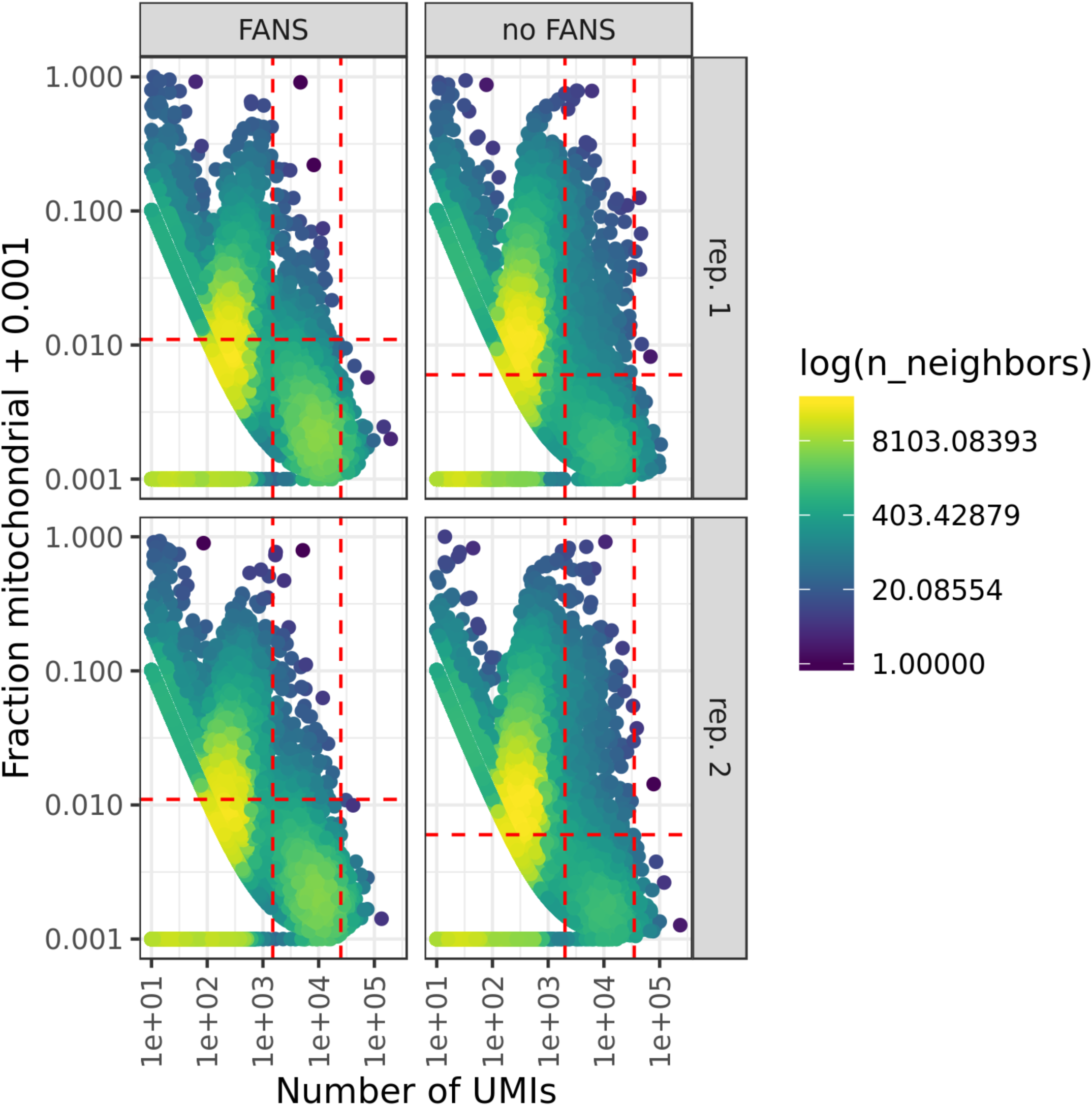
QC thresholds for FANS and non-FANS snRNA-seq libraries. Dashed lines represent thresholds for minimum number of UMIs, maximum number of UMIs, and maximum fraction of mitochondrial UMIs.

**Figure S4:**
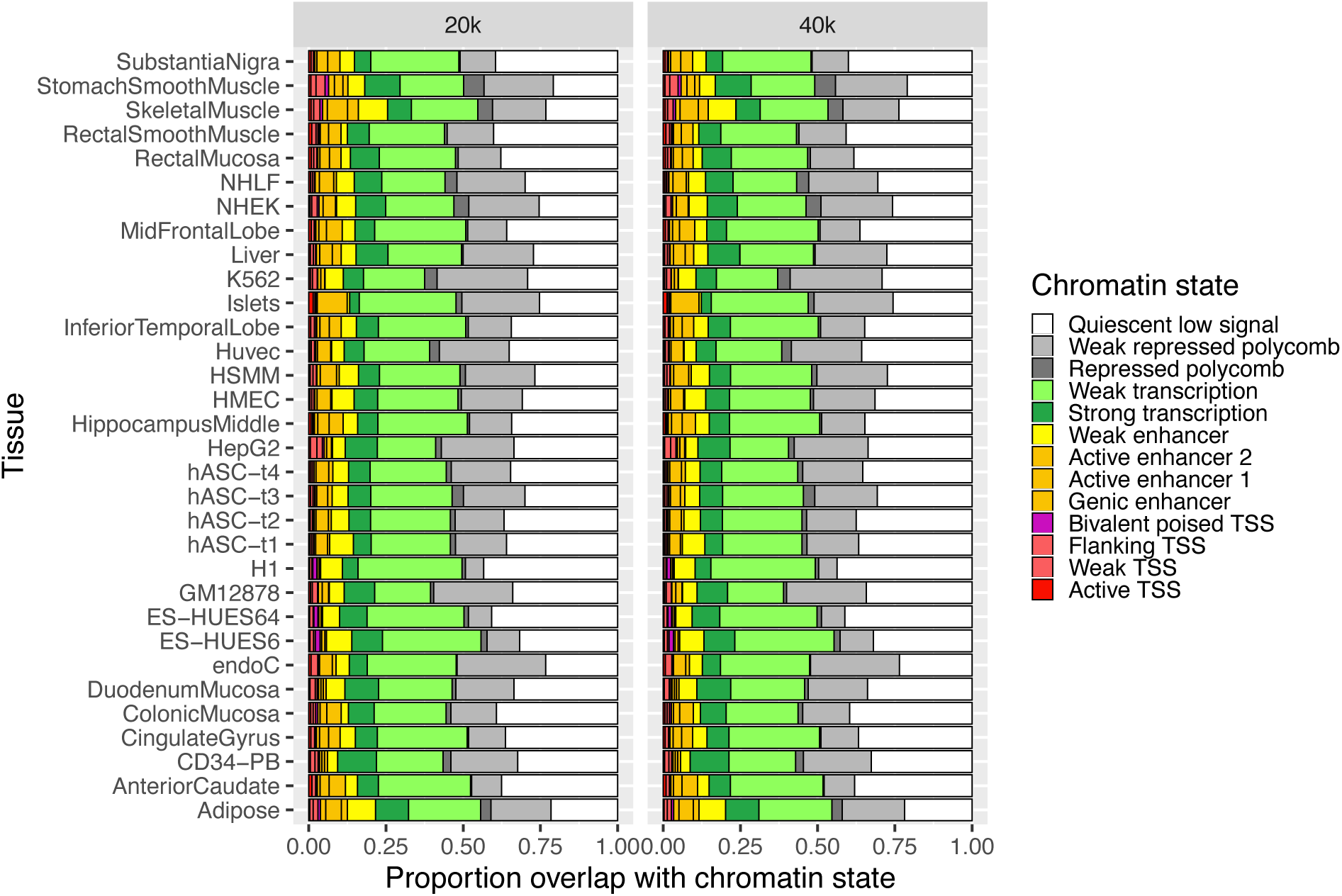
Chromatin state overlap for TSS-distal (>5 kb from TSS) ATAC-seq peaks from the 20k and 40k nucleus FANS snATAC-seq libraries.

**Figure S5:**
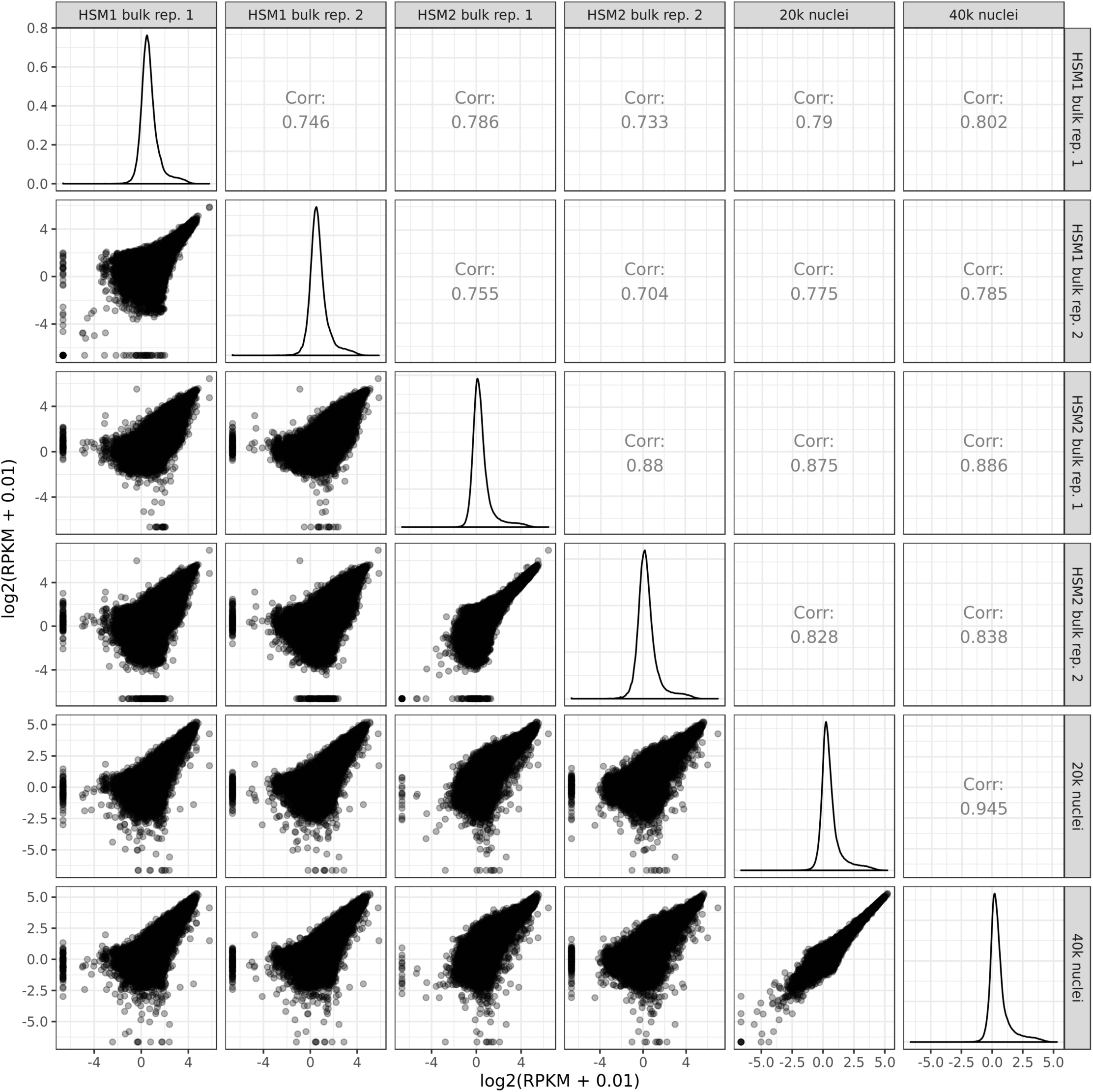
Correlation between 20k and 40k nucleus snATAC-seq libraries and standard bulk ATAC-seq libraries. Each point represents one peak.

**Figure S6:**
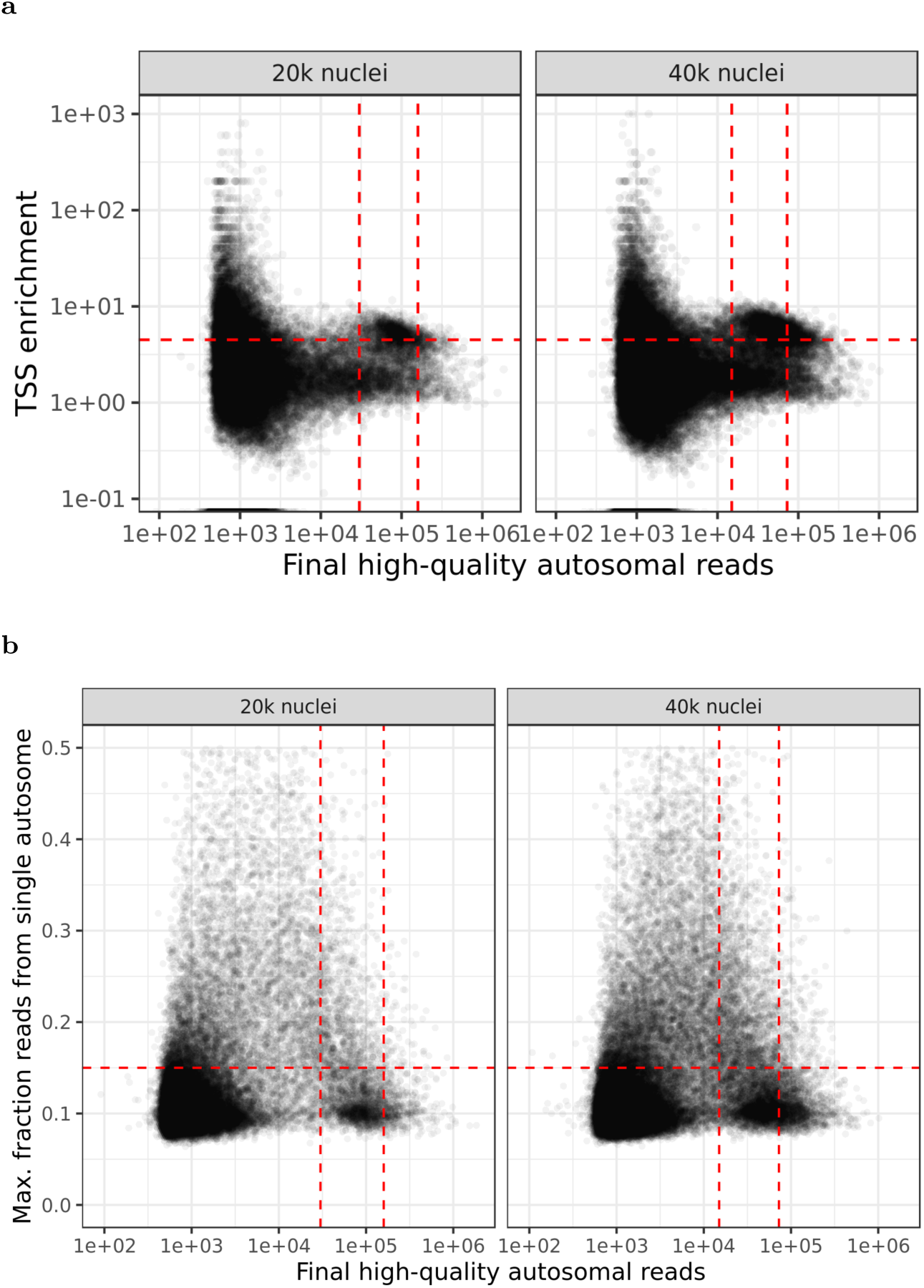
QC thresholding for the 20k and 40k nuclei input snATAC-seq libraries. (a) Dashed lines represent thresholds for minimum number of reads, maximum number of reads, and minimum TSS enrichment. (b) Dashed lines represent thresholds for minimum number of reads, maximum number of reads, and the maximum fraction of reads derived from a single autosome (imposed to filter out nuclei showing aberrant per-chromosome coverage).

**Figure S7:**
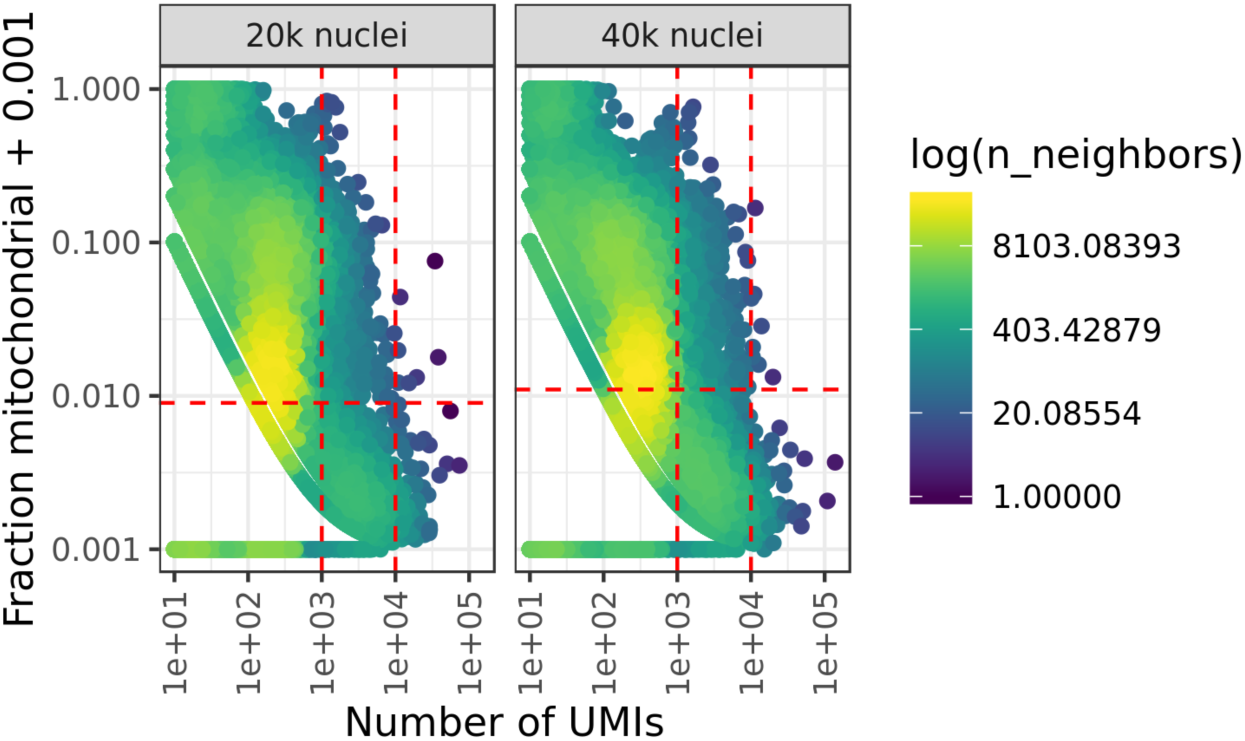
QC thresholds for the 20k and 40k nuclei input snRNA-seq libraries. Dashed lines represent thresholds for minimum number of UMIs, maximum number of UMIs, and maximum fraction of mitochondrial UMIs.

**Figure S8:**
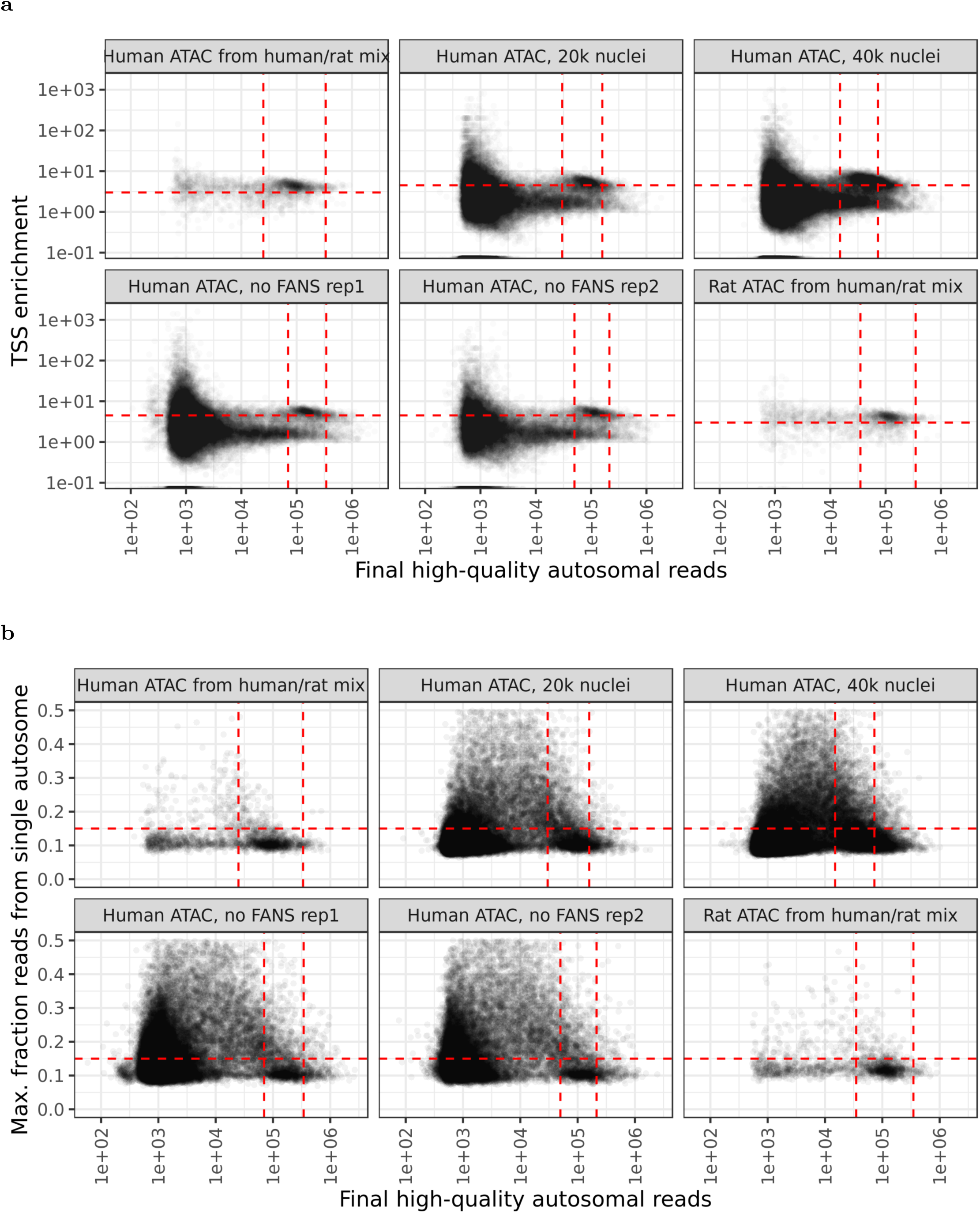
QC thresholds for all snATAC-seq libraries used in cell type clustering and downstream analyses. (a) Dashed lines represent thresholds for minimum number of reads, maximum number of reads, and minimum TSS enrichment. (b) Dashed lines represent thresholds for minimum number of reads, maximum number of reads, and the maximum fraction of reads derived from a single autosome (imposed to filter out nuclei showing aberrant per-chromosome coverage).

**Figure S9:**
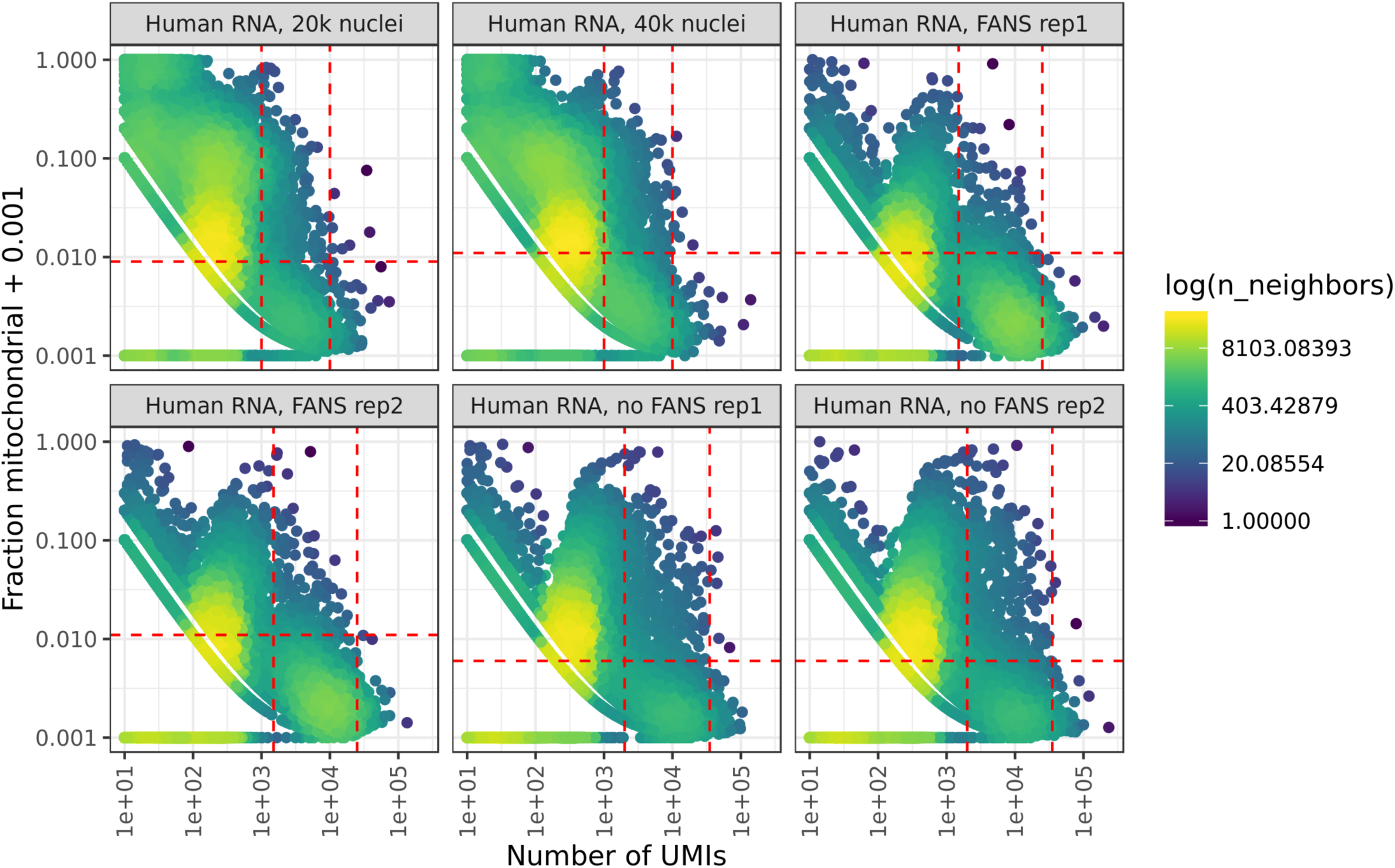
QC thresholds for all snRNA-seq libraries used in cell type clustering and downstream analyses. Dashed lines represent thresholds for minimum number of UMIs, maximum number of UMIs, and maximum fraction of mitochondrial UMIs.

**Figure S10:**
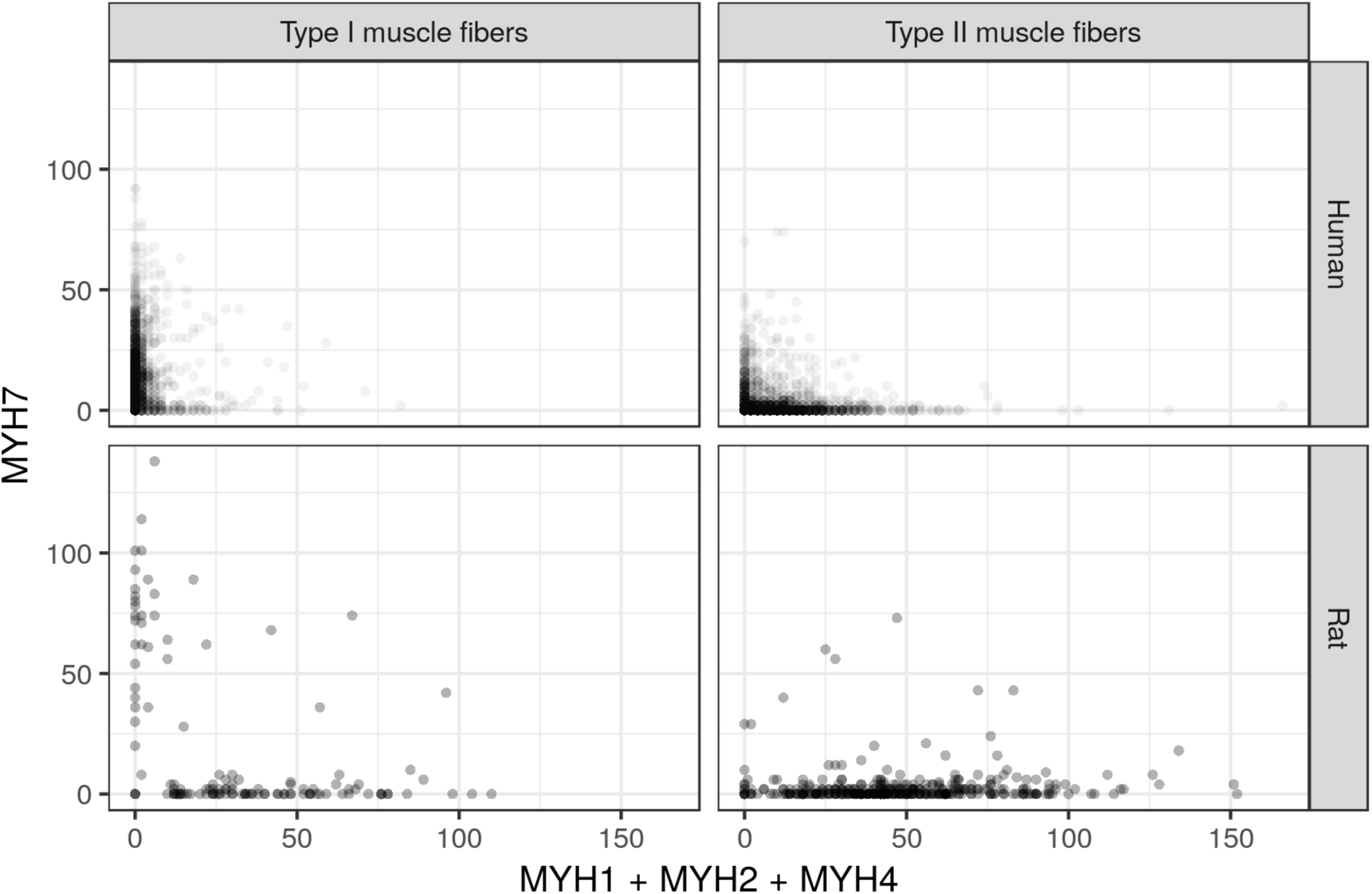
snATAC-seq read counts (gene promoter + gene body) derived from the Type II muscle fiber myosin heavy chain genes (MYH1, MYH2, MYH4) or the Type I muscle fiber myosin heavy chain gene (MYH7) for human and rat nuclei. Each point represents a single nucleus. Type I muscle fibers/Type II muscle fibers headers represent the cluster to which each nucleus was assigned.

**Figure S11:**
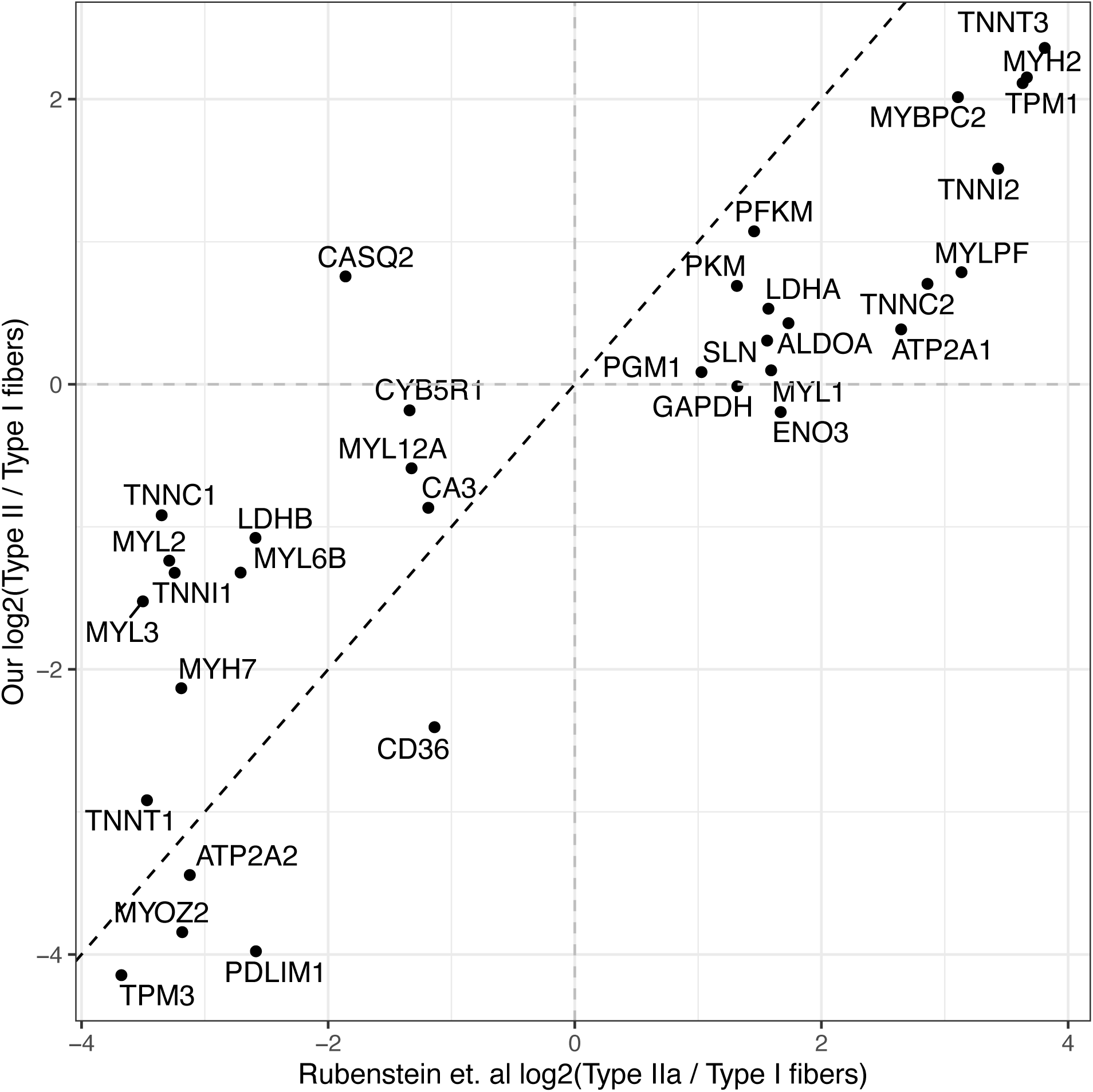
Log2(fold change) for Type II vs Type I muscle fiber gene expression, showing the genes with the largest fold changes between fiber types based on data from Rubenstein et al. (Rubenstein et al. Table S4). Rubenstein et al. performed RNA-seq on pooled type I and pooled type II muscle fibers, and determined the 20 genes with the largest fold change in type II relative to type I fibers, and the 20 genes with the largest fold change in the other direction, along with p-values for differential expression. The 34 genes (of those 40 genes) that were differentially expressed are shown here. The gene fold changes based on the muscle snRNA-seq data are often lower in magnitude than the fold changes based on Rubenstein et. al’s pooled RNA-seq data; this is unsurprising, as ambient RNA in the snRNA-seq data as well as any errors in nucleus fiber type assignments in snRNA-seq data clustering will reduce the observed fiber type differences.

**Figure S12:**
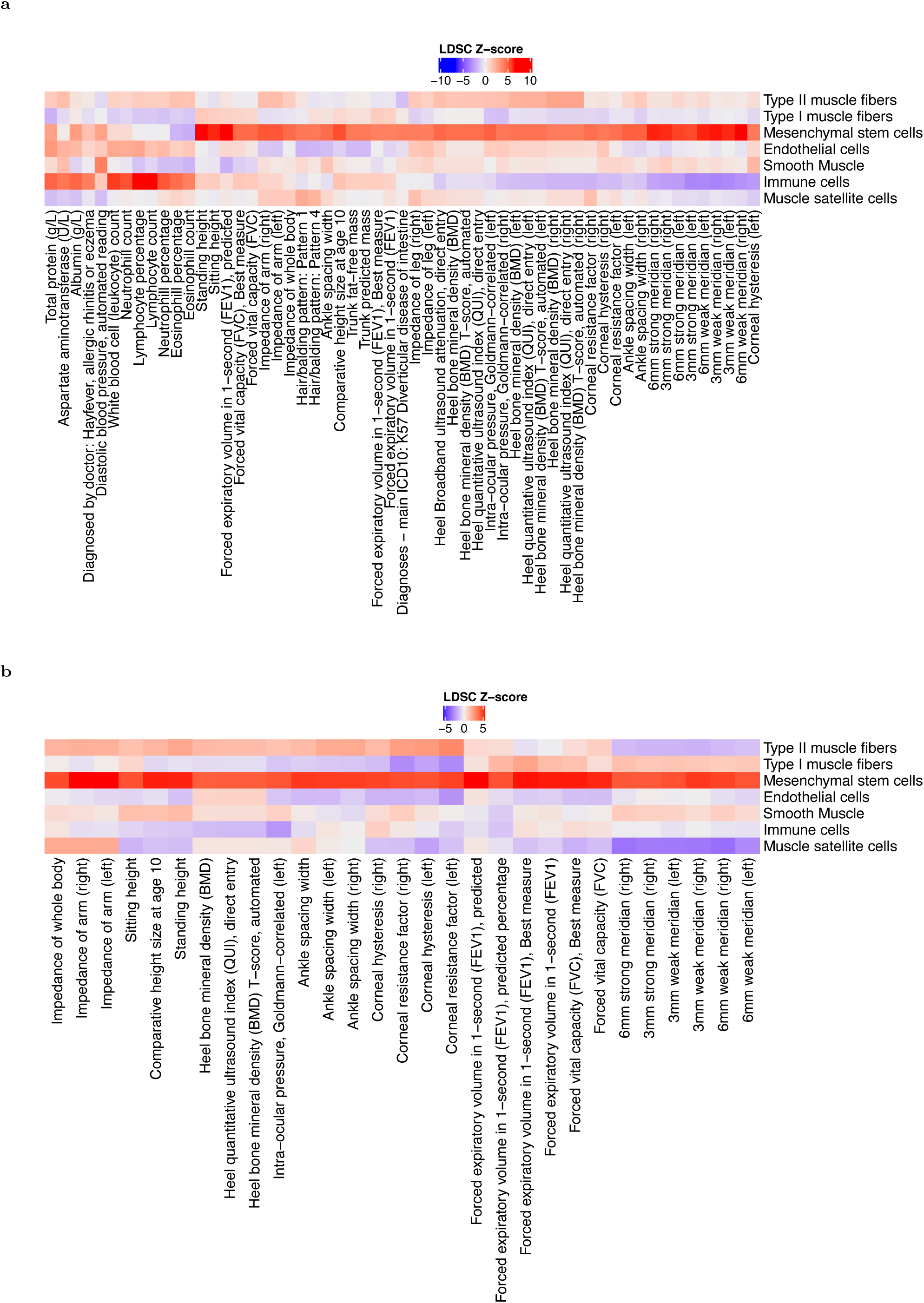
UK Biobank LDSC partitioned heritability results for traits for which one of the muscle cell types was significant after Benjamini-Yekutieli correction. (A) human, (B) rat.

**Figure S13:**
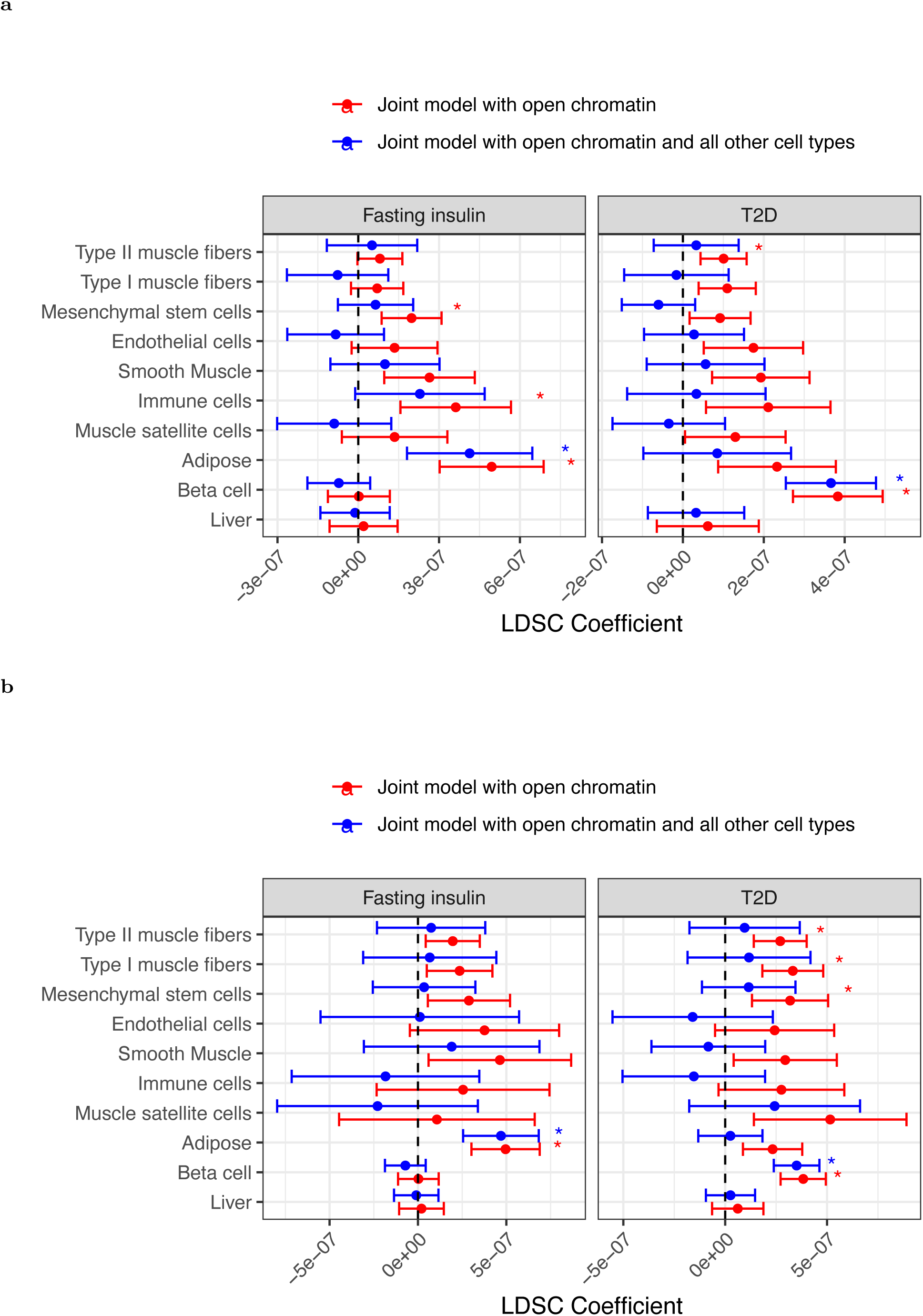
(A) LDSC partitioned heritability results for T2D (BMI-unadjusted) and Fasting insulin GWAS (BMI-adjusted), using human peak calls. Results are shown for pancreatic beta cell, adipose, and liver open chromatin regions as well. First, for each of the ten cell types, one model was run adjusting for cell type-agnostic annotations from the LDSC baseline model and common open chromatin regions (this is the joint model with open chromatin). Then, a single model containing those same annotations and all ten cell types was run (this is the joint model with open chromatin and all other cell types). Asterisk represents Bonferroni significance (p < 0.05 after adjusting for two traits, ten cell types, and two models per cell type = 40 tests). (B) Same as (A), but using the rat peak calls projected into human coordinates for the muscle cell types.

**Figure S14:**
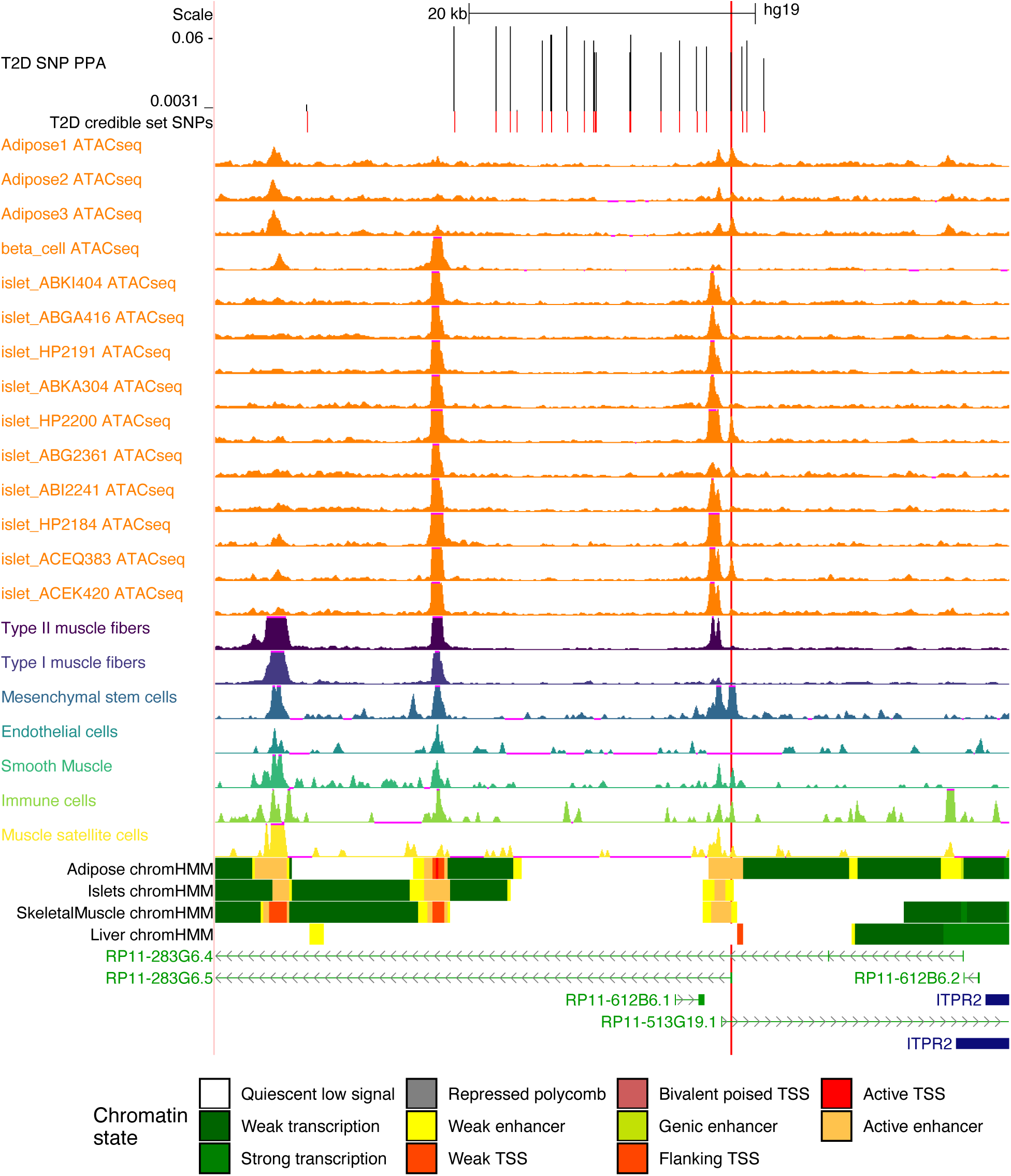
ATAC-seq signal in bulk adipose, bulk islet, single-nucleus pancreatic beta cell, or our muscle cell types at the *ITPR2* locus. Position of SNP rs7132434 is indicated by the long vertical red line. All tracks are normalized to 1M reads.

**Figure S15:**
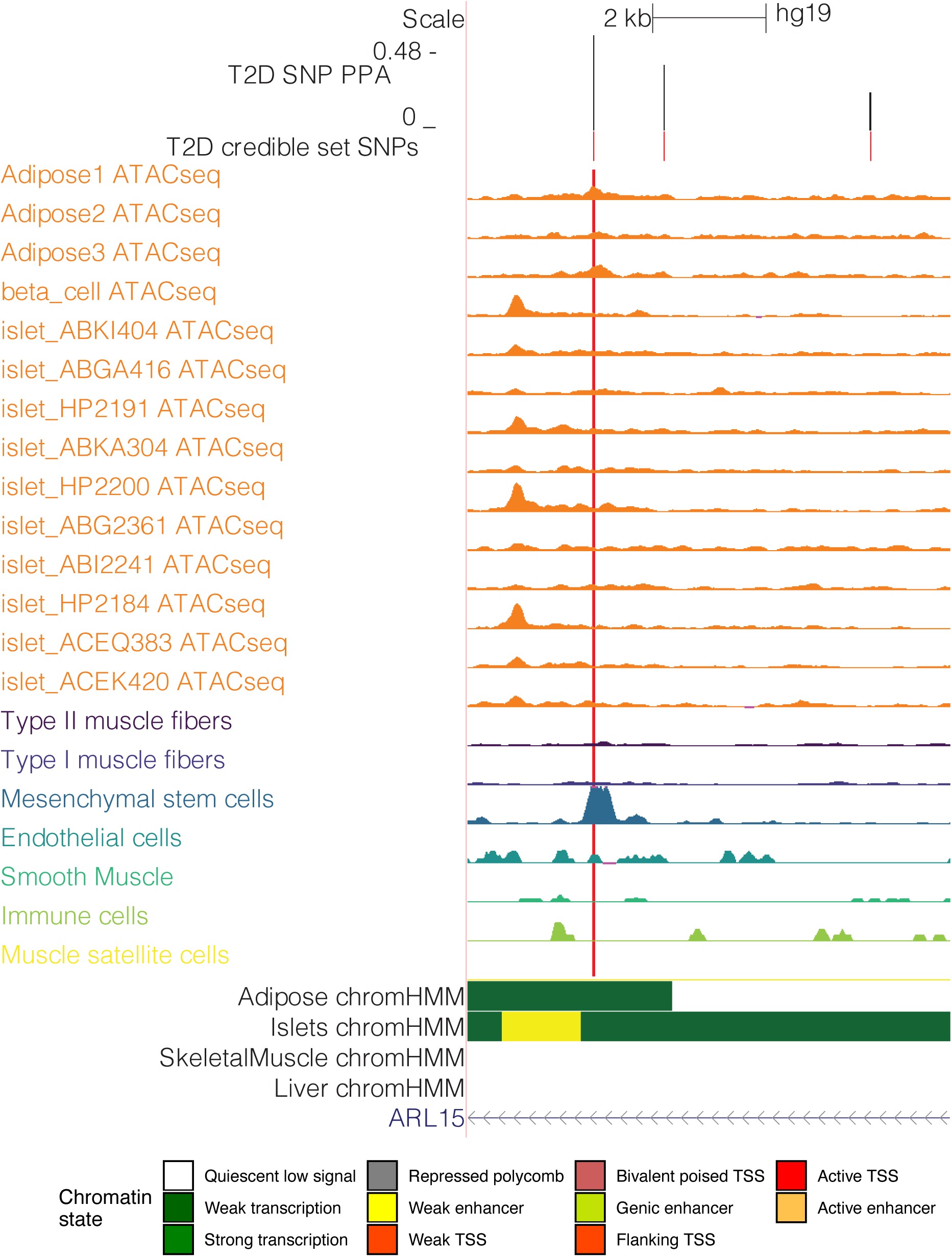
ATAC-seq signal in bulk adipose, bulk islet, single-nucleus pancreatic beta cell, or our muscle cell types at the *ARL15* locus. Position of SNP rs702634 is indicated by the long vertical red line. All tracks are normalized to 1M reads.

## Table legends

Table S1: snATAC-seq per-nucleus QC thresholds.

Table S2: snRNA-seq per-nucleus QC thresholds.

Table S3: Per library summary statistics (number nuclei per sample, mean and median fragments per library).

Table S4: Marker genes used for cluster cell type assignment.

Table S5: Nucleus counts per cell type, species, and modality.

Table S6: LDSC partitioned heritability z-scores for human snATAC-seq peaks.

Table S7: Cell type annotation overlap summary for credible sets from Mahajan et al. 2018. Values represent the number of credible set SNPs at each locus that overlap with the specified annotation. The *ARL15* locus discussed in the text (5_53271420) indicates one SNP overlaps with an Islet ATAC-seq peak; however, visual inspection of the locus reveals no convincing signal in any of the 10 examined islet ATAC-seq libraries (Fig. S15).

Table S8: LDSC baseline model annotations used.

