## Supplementary material for "Human and rat skeletal muscle single-nuclei multi-omic integrative analyses nominate causal cell types, regulatory elements, and SNPs for complex traits": Protocol S1

**Materials:**

- CP02 cryoPREP automated dry pulverizer (Covaris 500001)
- Eppendorf thermomixer C (EP 5382000015)
- 2mL glass tissue grinder and pestle (Kimble chase, 885301-0002, 8853000002)
- 40 µm strainer (Fisher, 22363547)

LB1 buffer should be made fresh each time.

| LB1 buffer | For 5mL | Final | Catalog number |
|---|---|---|---|
| 1 M HEPES, pH 7.5 | 0.25 mL | 50 mM | Invitrogen 15630080 |
| 5 M NaCl | 140 µL | 140 mM | Sigma S5150-1L |
| 0.5M EDTA, pH 8.0 | 10 µL | 1 mM | Promega V4231 |
| 50% glycerol | 1 mL | 10% | Sigma G5516-500 mL |
| NP-40 10% | 0.25 mL | 0.5% | Sigma 11332473001 |
| Triton X-100 10% (make 10% solution from the 100%) | 125 µL | 0.25% | Sigma T8787-100 mL |
| Ultra Pure Distilled water | 3.225 mL | | Invitrogen 10977015 |
| Immediately before use, add EDTA-free complete mini protease inhibitors (Roche). 1 tablet for every 5mL buffer. | 1 | | Roche 11836170001 |

**Samples: This protocol is designed for 4 samples. Please scale up or down based on the number of samples.**
**All the steps have to be performed on ice or at 4°C.**

**Nuclei extraction**

1. Frozen tissue (40-90 mg) was pulverized into fine powder while cold (dry ice and LN2) using an automated dry pulverizer CP02 cryoPREP.

2. Pulverized Frozen tissue (40-90 mg) was suspended in 1 mL of ice-cold 1x PBS in a 1.5 mL tube (Eppendorf 022431081) and centrifuged at 2000g for 3 min at 4°C. The supernatant was removed and the pellet was resuspended in 1 mL LB1.
3. The tissue was lysed by rocking the tubes in Eppendorf thermomixer C (EP 5382000015) at 4°C at 300 rpm for 10 min.
4. Each sample was transferred into a prechilled 2 mL glass Dounce homogenizer, and homogenized with 15 strokes of loose pestle A, and then transferred to 1.5 mL tube and centrifuged at 2000g for 5 min at 4°C.
5. The supernatant was aspirated and the pellet was resuspended in 1 mL of ice-cold 1 x PBS.
6. The suspension was filtered through 40 µm strainer and collected into a 6 well plate.
7. The well was washed with 1 mL of ice-cold 1 x PBS to collect all the remnant nuclei.
8. The nuclei were counted with cell counter (Trypan blue stains nuclei; typically 4-9 µm).
9. Nuclei were mixed and 154000 nuclei were transferred to 1.5mL tube, for each sample. If the volume was lower than 50µL, PBS was added to 1.5 mL tube to make the volume 50 µL.
10. The nuclei suspension was centrifuged again at 500 x g for 10 min at 4°C. The supernatant was aspirated very carefully (the supernatant was saved as a precaution) with flexi tip gel loading tip and the nuclei were resuspended in 20µL of diluted nuclei buffer to get a concentration of ~7700 nuclei/uL.
