## Supplementary material for "Human and rat skeletal muscle single-nuclei multi-omic integrative analyses nominate causal cell types, regulatory elements, and SNPs for complex traits": Protocol S2

**Materials:**

- CP02 cryoPREP automated dry pulverizer (Covaris 500001)
- Eppendorf thermomixer C (EP 5382000015)
- 2mL glass tissue grinder and pestle (Kimble chase, 885301-0002, 8853000002)
- 70 µm strainer (Fisher 501457900)
- Celltrics, 20 µm, 30µm cell strainer (Fisher NC9682496, Fisher NC9699018)
- Fisherbrand Sterile Plastic Culture (FACS) Tubes 149563C
- RNase Inhibitor (Thermofisher N8080119)
- Eppendorf Protein Lobind 1.5 mL tubes (Eppendorf 022431081)
- Eppendorf Protein Lobind 2.0 mL tubes (Eppendorf 022431102)

LB1 buffer should be made fresh each time.

| LB1 buffer | For 5mL | Final | Catalog number |
|---|---|---|---|
| 1 M HEPES, pH 7.5 | 0.25 mL | 50 mM | Invitrogen 15630080 |
| 5 M NaCl | 140 µL | 140 mM | Sigma S5150-1L |
| 0.5M EDTA, pH 8.0 | 10 µL | 1 mM | Promega V4231 |
| 50% glycerol | 1 mL | 10% | Sigma G5516-500 mL |
| NP-40 10% | 0.25 mL | 0.5% | Sigma 11332473001 |
| Triton X-100 10% (make 10% solution from the 100%) | 0.125 mL | 0.25% | Sigma T8787-100 mL |
| Ultra Pure Distilled water | 3.225 mL | | Invitrogen 10977015 |
| Immediately before use, add EDTA-free complete mini protease inhibitors (Roche). 1 tablet for every 5mL buffer. | 1 | | Roche 11836170001 |
| 1% BSA in PBS | For 10 mL | Final | Catalog number |

| | | | |
|---|---|---|---|
| BSA | 100 mg | 1% | Fisher NC0390268 |
| PBS | Q.S. to 10 ml | | Invitrogen 10010023 |

**Samples: This protocol is designed for 4 samples. Please scale up or down based on the number of samples.**

**All the steps have to be performed on ice or at 4°C.**

**All the 2 mL and 1.5 mL tubes used in this protocol are Protein LOBIND tubes from EPPENDORF.**

**Nuclei extraction:**

1. Frozen tissue (40-90 mg) was pulverized into fine powder while cold (dry ice and LN2) using an automated dry pulverizer CP02 cryoPREP.
2. Pulverized Frozen tissue (40-90 mg) was suspended in 1 mL of ice-cold 1x PBS in a 1.5 mL tube (Eppendorf 022431081) and centrifuged at 2000g for 3 min at 4°C. The supernatant was removed and the pellet was resuspended in 1 mL LB1.
3. The tissue was lysed by rocking the tubes in Eppendorf thermomixer C (EP 5382000015) at 4°C at 300 rpm for 5 min.
4. Each sample was transferred into a prechilled 2 mL glass dounce homogenizer, and homogenized with 10 strokes of loose pestle A, and 20 strokes of tight pestle B and then transferred to 1.5 mL tube and centrifuged at 2000g for 5 min at 4°C.
5. The supernatant was aspirated and the pellet was resuspended in 1 mL of ice cold 1% BSA and centrifuged at 100g for 1 min at 4°C.
6. The supernatant was collected, discarding the loose debris pellet.
7. All the filters were prewet; 70 µm, 30 µm and 20 µm filters, with 200 µL of 1% BSA each and previously collected supernatant was sequentially filtered through 70 µm, 30 µm and 20 µm filters respectively.
8. The supernatant was filtered through 70 µm strainer and the filtrate was collected into a 50 mL conical tube.
9. The collected filtrate was passed through 30 µm celltrix strainer and collected into a 2mL tube.
10. The collected filtrate was passed through 20 µm celltrix strainer and collected into a 2mL tube.
11. The filtrate was transferred to a 1.5mL tube, and centrifuged at 350 x g for 10 min at 4°C (be careful of the tube direction). The supernatant was aspirated (the supernatant was saved as a precaution) with flexi tip gel loading tip (were very careful not to disturb the pellet) and the nuclei was resuspended in 500 µL of 1% BSA in PBS.
12. The nuclei suspension was centrifuged again at 350 x g for 10 min at 4°C (be careful of the tube direction). The supernatant was aspirated very carefully (the supernatant was saved as a

precaution) with flexi tip gel loading tip and the nuclei was resuspended in 100µL of 1%BSA in PBS.

13. The nuclei were counted with cell counter (Trypan blue stains nuclei; typically 4-9 µm) and diluted appropriately for RNA and ATAC submissions.
14. **(RNA Submission** ) To achieve the desired nuclei concentration, appropriate amount of nuclei was diluted with 1% BSA in PBS. To this suspension RNase inhibitor was added to get a final concentration of 0.2 U/µL. The nuclei was counted and submitted for snRNA seq.
15. (**ATAC Submission**)- The rest of the nuclei was spun down at 350 x g for 10 min at 4°C (were careful of the tube direction). The supernatant was aspirated very carefully with flexi tip gel loading tip and the nuclei was resuspended in appropriate volume of 1X diluted nuclei buffer (20x buffer supplied by 10X Genomics). The nuclei was counted and submitted for snATAC seq.
