## Supplementary material for "Human and rat skeletal muscle single-nuclei multi-omic integrative analyses nominate causal cell types, regulatory elements, and SNPs for complex traits": Protocol S3

**Transposition reaction**

1. A master mix of the following reagents was made, then 25 µL master mix was added to each nuclei sample.

| Transposition Mix | 1x | |
|---|---|---|
| TD (2X reaction buffer) | 12.5 | µl |
| TDE1 (Tn5 Transposase) | 2.5 | µl |
| nuclease-free H2O | 10 | µl |

2. The samples were incubated at 37°C for 1hr on thermomixer with shaking at 300 rpm
3. Qiagen MinElute kit was used to isolate transposed DNA in 11 µL EB.

**PCR 1st amplification**

**Library Amplification**

4. Transposed DNA fragments were amplified by combining the following reagents in a PCR tube

| PCR Mix | 1x | |
|---|---|---|
| nuclease-free H2O | 12.5 | µl |
| NEBNext High-Fidelity 2× PCR Master Mix | 25 | µl |

5. 37.5 µl of the PCR mastermix was added to each 0.2 µL PCR tube containing 10 µL of transposed DNA
6. 2.5 µl of 25µM unique barcoded forward/ reverse primer and 10µl of transposed DNA were added and amplified as follows.

*This first 5-min extension at 72°C is critical to allow extension of both ends of the primer after transposition, thereby generating amplifiable fragments. This short pre-amplification step ensures that downstream quantitative PCR (qPCR) quantification will not change the complexity of the original library.*

(1) 72°C 5 min
(2) 98°C 30 sec
(3) 98°C 10 sec
(4) 63°C 30 sec
(5) 72°C 1 min
(6) Repeat steps 3-5, 4x
(7) Hold at 4°C

7. The qPCR master mix was prepared while the PCR was running. In order to reduce GC and size bias in PCR, for the first PCR reaction do 5 cycles as above. qPCR was used here to monitor how many more PCR cycles are needed to avoid saturation.
8. A master mix of the following reagents was made:

| qPCR Master Mix | 1x | |
|---|---|---|
| SYBR green 100x | 0.09 | µl |
| nuclease-free H2O | 3.66 | µl |
| NEBNext High-Fidelity 2× PCR Master Mix | 5 | µl |

11. 8.75 µl of qPCR master mix was added into each tube.
12. 1.25 µl of 10µM unique forward /reverse primer was added to each tube.
13. 5 µl of template DNA (transposed and amplified 5 cycles) was added.
14. qPCR was run with the following cycles:

| | |
|---|---|
| 1 cycle: | 30 sec 98°C |
| 30 cycles: | 10 sec 98°C |
| | 30 sec 63°C |
| | 1 min 72°C. |

15. The number of cycles to amplify each sample was calculated from the Rn values. The maximum Rn value for each sample was taken and multiplied by 1/3. The cycle at which this value occurred for each sample was noted.
16. The Cycle value at which 1/3 of max Rn occurred, was the number of cycles the library was further amplified in the below PCR
17. A constant number of amplification cycles for all samples was determined and all the samples were amplified accordingly
18. The remaining 45 µL of PCR reaction was put back into the PCR machine:
    (1) 98°C 30 sec
    (2) 98°C 10 sec
    (3) 63°C 30 sec
    (4) 72°C 1 min
    (5) Repeat steps 2-4, x times (the calculated constant number of amplification cycles was used, usually 2-8 cycles)
    (6) Hold at 4°C
19. Amplified library was purified using Qiagen MinElute PCR purification kit and eluted in 35 µL of EB.
20. 5µL of the library was sent to the Advanced Genomics Core to run the Bioanalyzer to check for periodicity
21. If the Bioanalyzer looked good, DNA concentration of the rest of the sample was determined using Qubit and the samples were submitted for next-generation sequencing.
